# Multifaceted conserved functions of Notch during post-embryonic neurogenesis in the annelid *Platynereis*

**DOI:** 10.1101/2025.03.09.642222

**Authors:** Loïc Bideau, Loeiza Baduel, Gabriel Krasovec, Caroline Dalle, Ombeline Lamer, Mélusine Nicolas, Alexandre Couëtoux, Corinne Blugeon, Louis Paré, Michel Vervoort, Pierre Kerner, Eve Gazave

**Author notes:** Correspondence to: Eve Gazave. Deceased (Michel Vervoort).

## Abstract

Notch signaling is an evolutionarily conserved pathway known to orchestrate neurogenesis by regulating the transition of progenitor cells to differentiated neurons and glia, as well as by directing neurite outgrowth and axon guidance in many species. Although extensively studied in vertebrates and ecdysozoans, the role of Notch in spiralians (including mollusks, annelids or flatworms) remains largely unexplored, limiting our understanding of its conserved functions across bilaterians. In this study, we focus on the segmented annelid *Platynereis dumerilii*, a model organism in neurobiology and regeneration, to investigate Notch signaling functions during post-embryonic developmental processes. We show that Notch pathway components are expressed in neurogenic territories during both posterior elongation and regeneration, two processes requiring sustained neurogenesis. Through chemical inhibitions of the pathway and RNA-seq profiling, we demonstrate that Notch signaling regulates neural progenitor specification, differentiation, and overall neurogenic balance in the regenerating and elongating posterior part. Moreover, disruption of Notch activity leads to severe defects in pygidial and central nervous system organization, including abnormal axon guidance and impaired neurite outgrowth. Altogether, our results support the hypothesis that Notch has multifaceted conserved functions in neurogenesis across bilaterians, shedding light on the ancestral functions of this critical pathway.

## INTRODUCTION

Notch is an ancient signaling pathway, probably already functional in the last common ancestor of Metazoa [1, 2], which modulates a large array of cell fate decisions in a variety of developmental processes [3, 4]. Notch is a membrane receptor that is cleaved upon binding the ligand Delta/Jagged, located on a neighboring cell. Thus, Notch pathway is a contact-dependent (or juxtacrine) signaling [1, 5–7]. After cleavage, its intracellular domain is translocated into the nucleus of the receiving cell where it interacts with a transcriptional complex that regulates a large number of genes, including members of the Hairy Enhancer of Split (or Hes) superfamily [8, 9].

Notch is renowned for its fine-tuned regulation of cellular transitions from a progenitor to a differentiated state [4, 10]. It especially regulates the balance between progenitor cells and differentiated cells through a process known as lateral inhibition [4, 11], whereby a cell inhibits its neighboring cells from adopting the same fate. This is especially crucial during neurogenesis, where lateral inhibition through Notch signaling is key for regulating the transition from neural progenitor cells to mature neuronal cells (*i.e.* neurons) and/or glial cells. In short, Notch signaling is activated among a cluster of proneural cells where it specifies a neural precursor cell to become either a glial or a neuronal cell. This process maintains an essential balance for proper nervous system development in different animals such as in *Drosophila* [4, 12, 13] or in vertebrates [14, 15]. In both lineages, the inhibition of Notch signaling triggers similar neurogenic phenotypes consisting in an excess of neural cells [14]. In addition, Notch regulates axon guidance and neurite outgrowth in both *Drosophila* [16–18] and vertebrates [19, 20] during embryonic neuronal maturation. As neurogenesis is a lifelong process, Notch signaling is also involved during homeostatic adult neurogenesis [15, 21]. While data from vertebrates and *Drosophila* may suggest conserved roles of Notch in neurogenesis, the scarcity of functional analyses in the 3^rd^ large bilaterian lineage, the spiralians, hinders the ability to draw definitive conclusions regarding the evolution of its functions at the bilaterian scale [2, 22].

Among spiralians, the segmented annelid *Platynereis dumerilii* has emerged as a powerful model for diverse research fields such as development, evolution, regeneration, and neurobiology [23, 24]. Its complex larval nervous system encompasses both a central nervous system (CNS) and a peripheral nervous system (PNS). CNS consists on an anterior brain, and a ventral midline separating a stratified neurectoderm and a *bona fide* ventral nerve cord (VNC). The peripheral nervous system is connected to the CNS and is composed by ganglions innervating the appendages (parapodia), and by many sensory structures (cirri, tentacles …). *Platynereis*’ larval trunk neurectoderm is organized through proliferative/differentiating apico-basal layers and medio-lateral domains, prefiguring longitudinal tracks of different neuron types. This organization highlights key evolutionary conserved processes, such as the neurectoderm medio-lateral patterning [25] or the conservation of proneural bHLH gene functions among bilaterians [26]. Previous studies identified the core members of Notch pathway as well as its putative target *Hes* genes [8, 27], constituting a complete Notch repertoire in *Platynereis.* These genes were shown to be involved during embryogenesis in the correct patterning of the structure producing extracellular bristles or chaetae, the chaetal sac cells, likely through a lateral inhibition mechanism. Yet, no major role during embryonic or early larval neurogenesis was observed [27]. This prompted us to investigate the function(s) of Notch pathway during two key post-embryonic developmental processes in *Platynereis* - posterior elongation and posterior regeneration - which both require a sustained neurogenesis to innervate the newly formed tissues [28, 29]. Juveniles elongate by the addition of newly formed segment at their posterior end, thanks to a growth zone (GZ) of active progenitor or stem cells [28]. Juveniles have also the ability to replace a lost or injured posterior part [30] through restorative regeneration [31, 32]. Similar to most of regeneration processes, *Platynereis’* posterior regeneration can be divided into three common and sequential steps [33]. First, a wound healing closes the wound and produces a wound epithelium. The second step usually relies on the formation of a regeneration-specific structure called a blastema, following the mobilization of precursor cells. Late morphogenetic processes involving patterning, differentiation and growth of the reformed structure constitute the third and final step [32–34].

Here, we showed that the core members of the Notch pathway are dynamically expressed in all neurogenic territories during all steps of regeneration as well as during posterior elongation. Thanks to chemical inhibition and differential RNA-seq analyses, we found that Notch, through the action of its ligand *Delta* and potentially several *Hes* genes, regulates both the neural progenitor determination and the differentiated neurons balance in the regenerated pygidium (corresponding to the most posterior part of the worm). During posterior elongation, Notch alteration disturbs the neurogenetic cascade dynamic inducing a thickened CNS neurectoderm and an abnormally-shaped VNC with axon guidance and neurite growth defects. Altogether, our study supports the idea that Notch plays multiple and conserved roles during adult neurogenesis in bilaterians.

## RESULTS

### Dynamic expression of Notch pathway components and target genes in neurogenic structures during *Platynereis’* posterior regeneration

We previously determined that *Platynereis*’ genome contains the core components of the Notch pathway, including receptor *Notch*, two canonical ligands (*Delta* and *Jagged*), a series of putative alternative ligands (*Delta/Serrate-like* or *DSL* genes), as well as the gamma-secretase machinery and the negative regulator *Nrarp* [27]. Additionally, we previously characterized the *Hes* gene superfamily, identifying 15 members as putative Notch targets [8, 27]. In order to evaluate the role(s) of the Notch pathway in the context of posterior regeneration in *Platynereis,* we first analyzed the expression of its ligands, receptor, regulator and downstream targets using our previously generated RNA-seq data (Fig. 1A, Supp. Fig. 1A, Supp. Table 1) [35]. Most Notch-related genes displayed dynamic expression, with upregulation peaking at later regenerative stages (from 3 dpa – days post-amputation - onward), suggesting a role in blastema growth and differentiation (Fig. 1A, Supp. Fig. 1A, Supp. Table 1).

**Figure 1:**
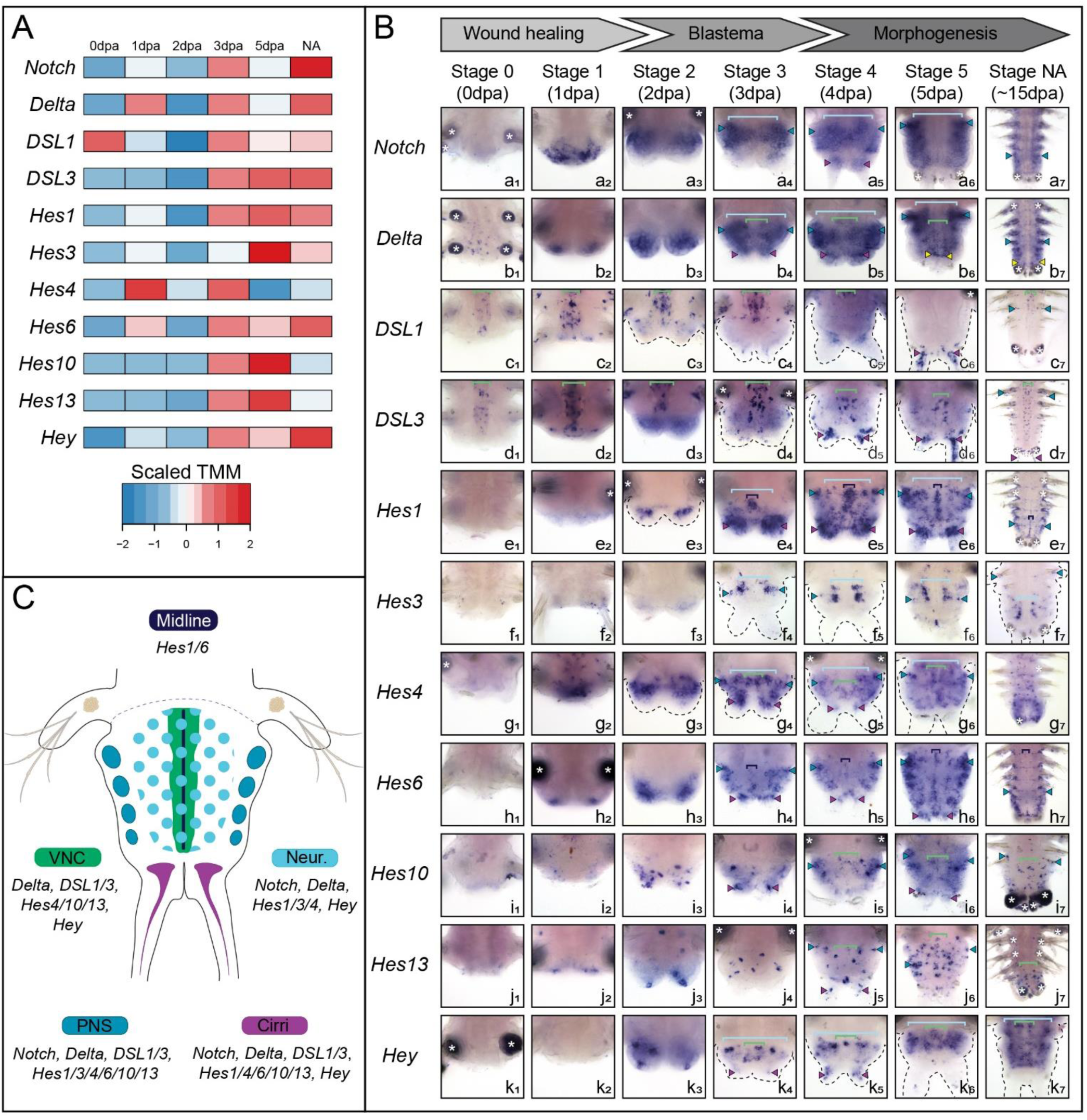
Several core members of the Notch pathway and its putative target genes are dynamically expressed in neurogenic structures during posterior regeneration. A) Heatmap representation of expression levels of several Notch components and *Hes* genes during posterior regeneration [35]. B) Whole-mount *in situ* hybridizations (ventral views) of Notch components and *Hes* genes expressed in a neurogenic structure during three main steps of regenerative processes (wound healing, blastema formation and morphogenesis). C) Schematic drawing of a regenerated part at stage 5 depicting the expression patterns of the genes found expressed in a neurogenic structure during regeneration. VNC = ventral nerve cord (green brackets); PNS = peripheral nervous system (blue arrowheads); Neur. = neurectoderm (light blue brackets); Cirri (purple arrowheads); Midline (dark blue brackets); yellow arrowheads = growth zone involved in posterior elongation of the animals [28]; white asterisks = non-specific staining from parapodial glands. dpa = day(s) post-amputation, NA = non-amputated.

We then assessed their expression patterns during posterior regeneration using whole mount *in situ* hybridizations (WMISH) thanks to an established staging system covering all main regeneration steps: wound healing, blastema formation and morphogenesis [29] (Fig. 1, Supp. Fig. 1). Strikingly, we observed that among the six core members of the Notch pathway, four of them are expressed in one or more neurogenic territories *i.e.* neurectoderm, ventral nerve cord (VNC), peripheral nervous system (PNS) and sensory anal cirri. Similarly, seven of the twelve *Hes*-related genes are expressed in a nervous system-related structures (Fig. 1B, C).

*Wound healing:* Immediately after amputation, most of the genes are not detected in the tissues proximate to the wound with the noticeable exceptions of *DSL1 & 3*, expressed in the VNC of the non-amputated (NA) tissues (Fig. 1B, stage 0 – green square brackets). At 1 dpa, the receptor *Notch* is widely expressed in the wound epithelium, while the ligands *Delta, DSL1* and *DLS3* and some *Hes* genes (*Hes4, 6, 13)* have a more discrete expression in few epithelial cells (Fig. 1B, stage 1).

*Blastema formation*: At 2 and 3 dpa, *Notch* and *Delta* are broadly expressed in both the mesoderm and the ectoderm, potentially the neurectoderm and the PNS of the small blastema (Fig. 1B, stages 2 & 3– light blue square brackets and blue arrowheads, respectively). *DSL1* and *3* are mainly restricted to the VNC of the NA tissues at 2 dpa (Fig. 1B, stage 2 – green square brackets). Conversely, *DSL3* expression extends to nerve cells from the newly regenerated VNC at 3 dpa (Fig. 1B, stage 3– green square brackets). At 2 dpa, some *Hes* genes are expressed in modest (*Hes1, 6*) to large areas (*Hes4*) in the ectoderm of the blastema (Fig. 1B, stage 2). At 3 dpa, these genes exhibit a strong expression in the midline (*Hes1, 6* - dark blue square brackets), cirri (*Hes1, 4, 6, 10, Hey* - purple arrowheads), neurectoderm (*Hes1, 3, 4, Hey* – light blue square brackets) and the PNS (*Hes3, 4,* 6 –blue arrowheads) (Fig. 1B, stage 3). A number of other genes are expressed in a restricted manner in few scattered ectodermal cells (*Hes10, 13*, *Hey*), which may be neural cells of the VNC, PNS and/or cirri at stages 2 and 3.

*Morphogenesis*: At 4, 5 and at 15 dpa (a proxy of NA posterior part [28]), all the genes exhibit broadly similar yet larger expression patterns than the ones observed at stage 3. *Notch* and *Delta* are largely expressed in mesodermal and ectodermal tissues including the neurectoderm, the VNC, the PNS and the anal cirri (Fig. 1B, stages 4, 5, NA – light blue square brackets, green square brackets, blue arrowheads and purple arrowheads, respectively). *DSL1* & *3* are expressed in neurons of the VNC and anal cirri (green square brackets and purple arrowheads respectively). *Hes* genes are found in all neural structures: the midline (*Hes1, 6 -* dark blue square brackets), the VNC (*Hes4, 10, 13, Hey -* green square brackets), the neurectoderm (*Hes1, 3, 4, Hey -* light blue square brackets), the PNS (*Hes1, 4, 6, 10, 13 -* blue arrowheads), and the cirri (*Hes1, 4, 6, 10*, *13, Hey* - purple arrowheads). Hence, a combination of at least the *Notch* receptor, one ligand and several *Hes*-related genes are found co-expressed in all neurogenic structures during regeneration (Fig. 1C).

However, the expression of Notch pathway members and *Hes* putative target genes expression is not restricted to neural territories. Instead, expression is observed in segmental stripes (*DSL2* and *Hes5*, Supp. Fig. 1B-a, d), the ventral vessel (*DSL2*, Supp. Fig. 1B-a), and a broad mesodermal area (*Hes8* and *11*, Supp. Fig. 1B-e, f). In addition, three genes (*Nrarp, Hes2* and *Hes 12)*, are expressed in chaetal sacs (Supp. Fig. 1B-b, c, g, pink arrowheads), the structures responsible for the production of chaetae composed of follicle cells surrounding a central chaetoblast (as previously described during larval development [8, 27]). Interestingly, *Delta*, *Nrarp*, *Hes2* and *Hes8* are expressed in the regenerated growth zone or GZ (Fig. 1B-b and Supp. Fig. 1Bb, c, e - yellow arrowheads) [28].

By combining RNA-seq data with the thorough analysis of complex and dynamic expression patterns of the Notch pathway components and putative *Hes* target genes, we hypothesize that Notch orchestrates – albeit not exclusively – key neurogenic functions during posterior regeneration and elongation in *Platynereis*.

### Early Notch inhibition alters neurogenesis and induces pygidial hypertrophy during *Platynereis’* posterior regeneration

We then performed a series of chemical inhibition experiments to determine the role(s) of Notch signaling during the 5-day process of posterior regeneration in *Platynereis*. We observed that as early as 2 dpa, the treated worms (exposed to all the tested gamma-secretase inhibitors - LY-411575, RO-4929097 and DAPT) displayed a statistically significant regeneration delay compared to DMSO controls, failing to progress beyond stage 2 (characterized by a small bi-lobed blastema) (Fig. 2A, Supp. Fig. 2A-C). To determine whether this delay resulted from impaired cell proliferation, a process known to be mandatory for regeneration to proceed [29], we performed a 1-h EdU pulse experiment in both DMSO control and LY-411575-treated worms. At 2 and 5 dpa, the proportion and distribution of EdU+ cells were comparable between conditions, indicating that proliferation remained unaffected (Fig. 2B, C). Additionally, a 3-day EdU chase starting at 2 dpa revealed extensive signal dilution in both control and treated worms, confirming ongoing cell divisions (Supp. Fig. 2D). Thus, the regenerative arrest observed upon Notch inhibition is not due to reduced proliferation. Therefore, given the limited growth of the regenerated structure, we assessed cell death profile with TUNEL assay (Fig. 2D, E). In DMSO controls, apoptotic cells were detected in internal tissues at 2 dpa, consistent with the expected response to amputation (unpublished lab data), but were nearly absent by 5 dpa. In contrast, LY-411575-treated worms exhibited a significantly higher proportion of TUNEL+ cells at 5 dpa, affecting both internal and superficial tissues (Fig. 2D, E). Thus, Notch inhibition increases cell death during posterior regeneration.

**Figure 2:**
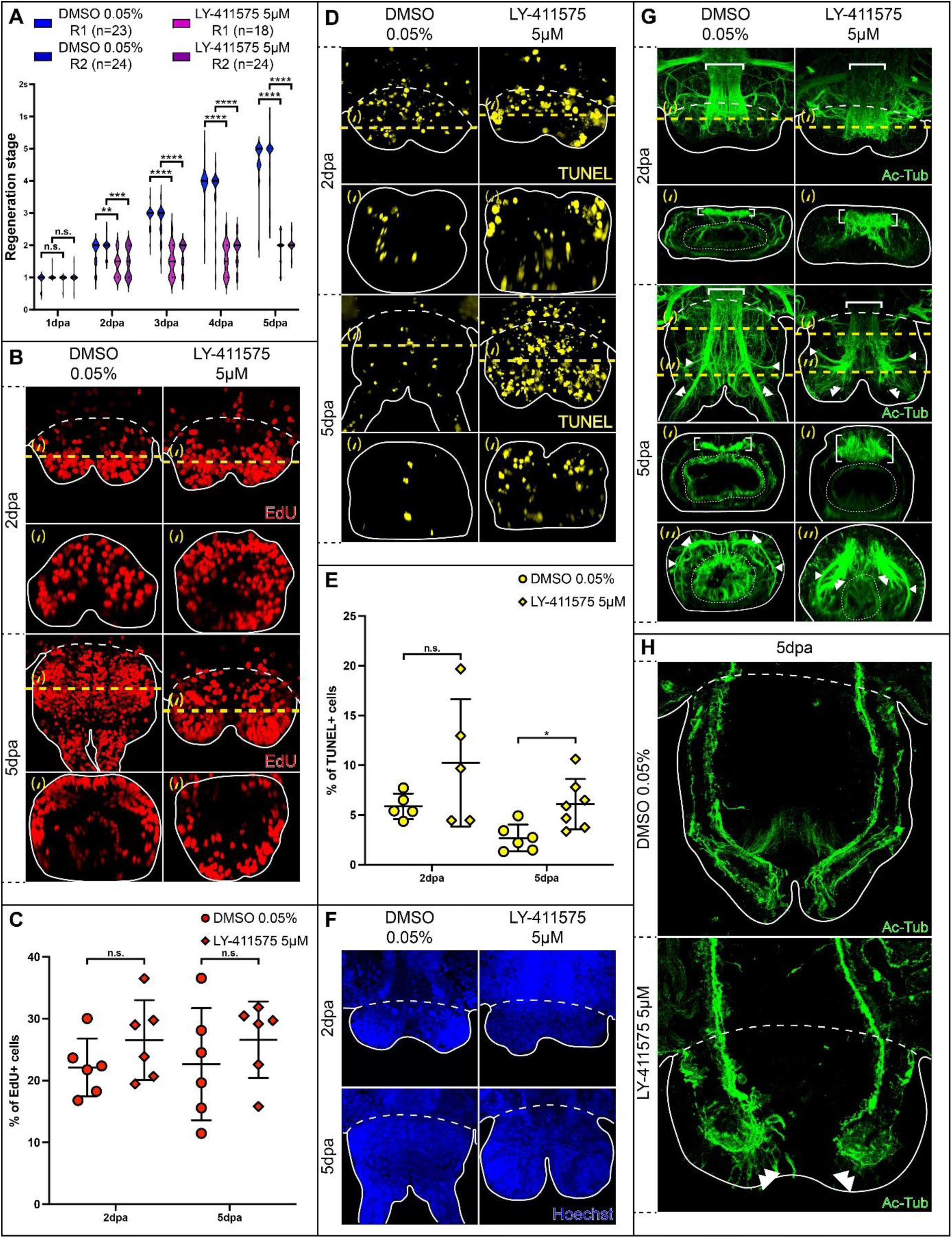
Morphological and cellular effects of Notch inhibition during posterior regeneration. A) Violin plots representing the regeneration stages reached by each animal every day for 5 days upon LY-411575 5 µM treatment in comparison to DMSO 0.05% control. R1 and R2 are independent replicates performed for n animals. B - F) EdU labelling following a 1h-pulse (B) and comparison of the proportions of EdU+ cells between conditions at 2 and 5 dpa (C); TUNEL assay (D) and comparison of the proportions of TUNEL+ cells (E); Hoechst nuclei labelling (F). G - H) Acetylated tubulin (Ac-tub) immunolabelling on whole-mount regenerated parts of LY-411575-treated worms and controls at 2 and 5 dpa. H) Acetylated tubulin (Ac-Tub) immunolabelling on longitudinal cross-sections of LY-411575-treated worms and controls at 5dpa. (B-D-G) Ventral views are on top and corresponding virtual transverse sections (along the yellow dotted lines (‘) or (‘’)) are at the bottom. In all relevant panels, solid white lines delineate the outlines of the samples, white dashed lines correspond to the amputation planes and white dotted lines delineate the gut. Each bracket corresponds to a Mann-Whitney U test. n.s. p>0.05; * p<0.05. dpa = day(s) post-amputation. White brackets = ventral nerve chord; white arrowheads = circular pygidial nerve; white double arrowheads = nerves of the anal cirri.

Detailed examination of the morphology and cell nuclei arrangements revealed that regenerated parts from LY-411575-treated worms are not merely arrested at approximately stage 2, but rather exhibits morphological defects. At 5 dpa, the regenerative structure appears hypertrophied, with enlarged blastemal lobes (Fig. 2F), even though their cellular densities remain unchanged (Supp. Figure 2E). To further characterize this phenotype, we conducted an extensive molecular analysis using a set of markers to label different structures and tissues involved in *Platynereis’* posterior regeneration (Fig. 2, Supp. Fig. 3) [28, 29, 36]. Ring-like expression of *Hox3* reveals that the ectodermal GZ is maintained upon Notch inhibition (Supp. Fig. 3A – yellow arrowheads) but is positioned more anteriorly in the 5dpa LY-411575-treated regenerated part (Supp. Fig. 3A’2) than in the control (Supp. Fig. 3A’1). Similarly, the expression patterns of *Evx, PiwiB, Myc* and *Nanos*, confirm the presence of a mesodermal GZ in treated worms (Supp. Fig. 3B-E – yellow arrowheads). However, these markers also indicate a significant reduction in the territories occupied by mesodermal progenitors within the blastema (Supp. Fig. 3B-E). Moreover, the typical striped expressions of the segmentation genes *Engrailed (en)* and *Wnt 1* [37], (Supp. Fig. 3F-G) are lost in LY-411575-treated worms at 5 dpa (Supp. Fig. 3F’2, G’2). In contrast, the expression patterns of markers of the terminal part of the worm, *i.e.* the pygidium and its pygidial cirri (*Cdx* and *Dlx*, Supp. Fig. 3H, I) are expanded, suggesting that most cells of the regenerating part at 5 dpa have a pygidial identity upon Notch inhibition. Unaltered expression patterns of *Fox A* (Supp. Fig. 3G, J) together with the presence of an enlarged ring-like bundles of pygidial muscles, shown by the expression of *Twist* and *TroponinI* (Supp. Fig. 3K, L) and phalloidin labelling (Supp. Fig. 3M, N) indicate proper gut and muscle regeneration within the hypertrophied structure. Finally, acetylated tubulin labelling of neurites revealed striking nervous system defects in this hypertrophied pygidium (Fig. 2G, H). While the circular pygidial nerve is preserved (white arrowheads), the two nerves innervating the anal cirri (white double arrowheads) are disrupted and multiple aberrant nerve projections extend throughout the 5 dpa regenerated structure (Fig. 2G, H). In addition, at both 2 and 5 dpa, the VNC (white brackets) exhibit a dimmer signal and is noticeably thicker in LY-411575-treated worms.

Taken together, these results indicate that early chemical inhibition of Notch pathway during posterior regeneration in *Platynereis* leads to the formation of an altered terminal structure, thereby hindering regeneration to proceed properly. This structure harbors a dysfunctional growth zone producing few tissues and a hypertrophic proliferative pygidium exhibiting increased apoptosis and severe nervous system disruptions.

### Notch signaling controls pygidial neurogenesis during regeneration by regulating *Hes* genes activity

#### Transcriptome-wide analysis of Notch inhibition identifies nervous system genes as downstream targets

Given the dramatic nervous system defects induced by Notch inhibition during posterior regeneration, we decided to further explore its impact on gene expression thanks to a bulk RNA-seq unbiased approach between LY-411575-treated and control worms at 1 and 2 dpa (Sup. Fig. 4A and B, Supp. Table 2).

Differential gene expression analysis indicated that at 1 dpa, 932 differentially expressed genes (DEG) were identified between DMSO control and LY-411575 treated worms. Among them, 401 are downregulated, 531 are up regulated and 313 are specific to this comparison (Fig. 3A, A’, Supp. Table 3). A similar number of DEG were identified between the control and treated worms at 2 dpa (n=1012; 439 are downregulated, 473 are up regulated and 356 are specific to this comparison) (Fig. 3A, A’, Supp. Table 4). Gene Ontology (GO) term enrichment analysis indicate that many genes related to stress are upregulated at both 1 and 2 dpa, while genes related to neurogenesis and development are downregulated (Supp. Fig. 5). The thorough manual analysis of the DEG at 1 and 2 dpa (both up and down combined) confirmed and extended this tendency. Genes related to inflammation, immune system, redox signaling and stress, processes well known to be involved in the early steps of regeneration [32, 38–40] are prominent. Importantly, one fifth of DEG are related to nervous system, more specifically to axon connections or neurite outgrowths regulation, consistent with the acetylated tubulin aberrant phenotype (Fig. 2G, H). Key members of the neurogenic cascade, such as *achaete-scute* 1 and 2 (markers of neural progenitors) and *atonal* (a marker of PNS neurogenesis) are upregulated more specifically at 2 dpa [26].

**Figure 3:**
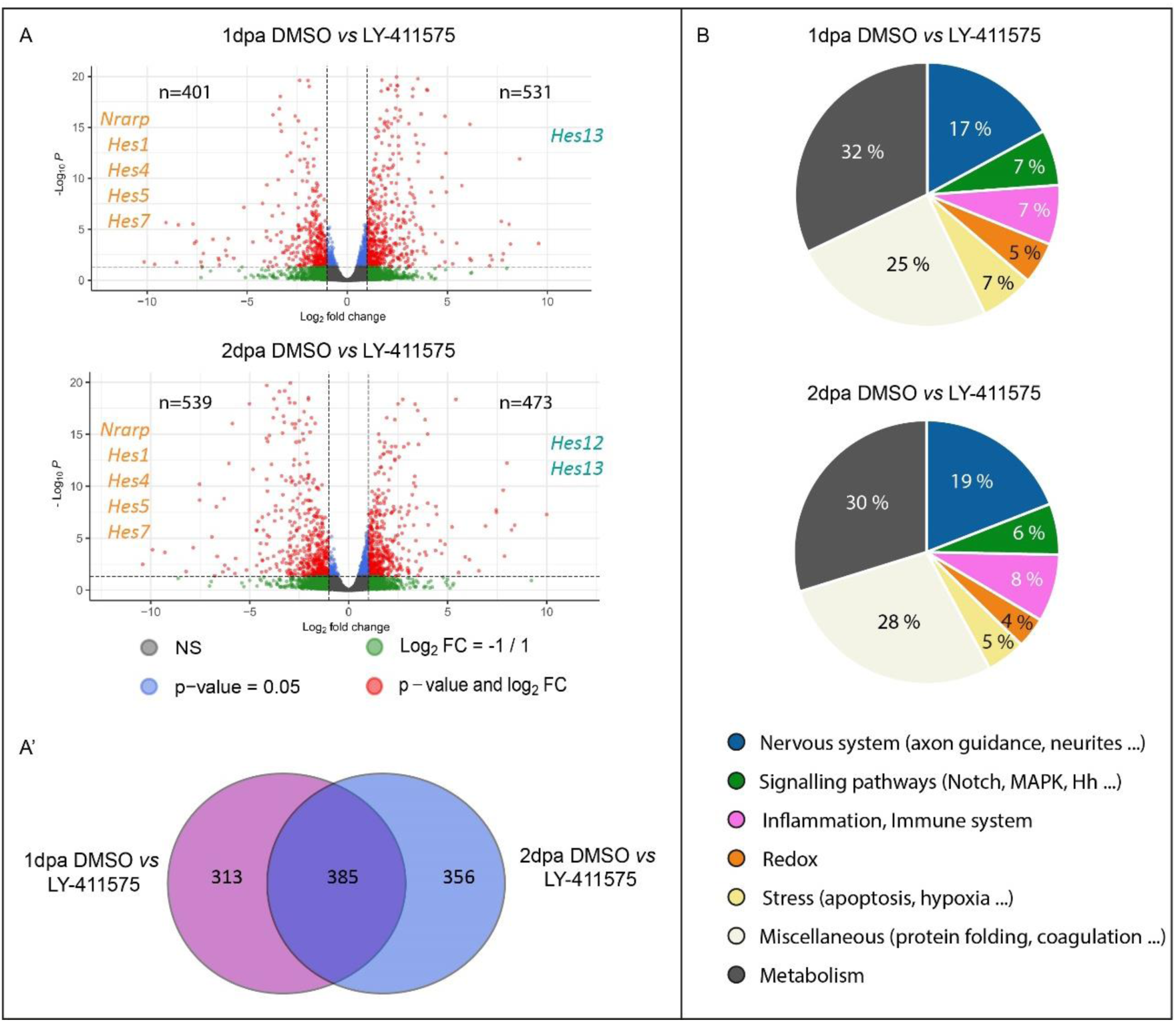
Impacts of Notch inhibition on transcriptome-wide gene expression during regeneration. A) Volcano plots comparing LY-411575-treated regenerated parts *vs* controls at 1 (top) and 2 dpa (bottom). Each dot represents a transcript, with colors indicating different values for FDR and logFC. In grey: FDR > 0.05; −1 < logFC < 1. In green: FDR > 0.05; logFC < −1 or logFC > 1. In blue: FDR < 0.05; −1 < logFC < 1. In red: FDR <0.05, −1 < logFC < 1. Thus, differentially expressed genes (DEGs) are depicted in red. Few examples of genes related to the Notch pathway upregulated in LY-411575-treated conditions are shown in cyan on the right of the plots while examples of downregulated genes are shown in orange on the left. A’) Venn diagram representing the DEGs specific or common among 1 and 2 dpa conditions. B) Pie charts depicting the functional classifications of the DEGs for 1 and 2 dpa conditions. dpa = day(s) post-amputation.

#### Notch inhibition triggers excessive neurogenesis and ectopic neurons formation in the hypertrophied regenerated pygidium

Using a set of markers known to be involved in the formation of the larval nervous system [25, 26, 41, 42], we assessed the effects of Notch inhibition on the developmental neurogenic cascade during pygidium regeneration (Fig. 4).

**Figure 4:**
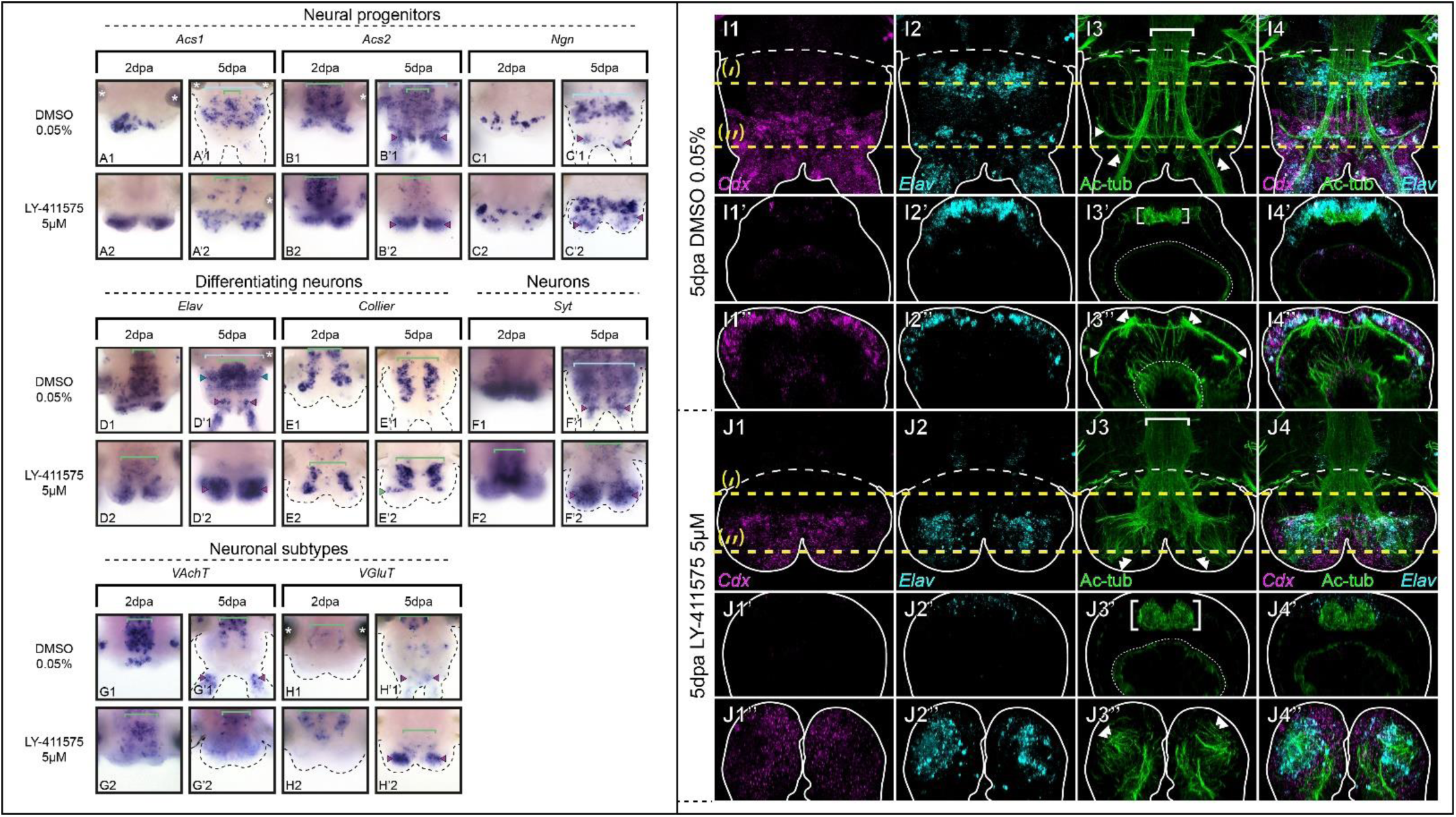
Notch signaling pathway inhibition during posterior regeneration leads to major neural defects in the pygidium. A to H) Whole-mount *in situ* hybridizations for markers of neural progenitors (A-C), differentiating neurons (D, E), differentiated neurons (F) and neuronal subtypes (G, H) for LY-411575 treated worms and controls at 2 and 5dpa. Ventral views. Green brackets = ventral nerve cord (VNC); light blue brackets = neurectoderm; purple arrowheads = neurons of the pygidium and cirri; green arrowheads = ectopic VNC neurons; white asterisks = non-specific staining from glands. (I-J) Hybridization chain reaction (HCR) for *Cdx* (purple, 1 and 4) and *Elav* (cyan, 2 and 4) coupled with immunolabelling for acetylated tubulin (green, 3 and 4) for LY-411575 treated worms and controls at 5dpa. Ventral views are on top and corresponding virtual transverse sections along the yellow dotted lines (‘ and ‘’, respectively) are at the bottom. Solid white lines delineate the outlines of the samples, white dashed lines correspond to the amputation planes. White brackets = VNC; white arrowheads = circular pygidial nerve; white double arrowheads = cirri nerves. dpa = day(s) post-amputation.

First, Notch inhibition altered the expression patterns of several neural progenitor markers (*Achaete-scute 1* and *2 (Acs1&2), Neurogenin (Ngn)*). At 2 dpa, *Acs1&2* were markedly upregulated in the blastema of treated worms, (Fig. 4A2, B2), whereas *Ngn* expression remained unchanged compared to the control (Fig. 4C2). At 5 dpa, these three markers were broadly expressed in the hypertrophied pygidium in LY-411575-treated worms (Fig. 4A’2, B’2, C’2 - purple arrowheads), in stark contrast to their restricted expression in controls (in particular for *Acs1* and *Ngn*, Fig. 4A’1, C’1 - purple arrowheads). Notably, *Acs1&2* were strongly downregulated in the neurectoderm of treated worms (Fig. 4A’2, B’2 – light blue square brackets), likely due to the severe reduction in tissue growth. We then determined the expression patterns of two markers of differentiating neurons: *Elav*, found in all differentiating neurons (Fig. 4D, VNC – green square brackets, PNS – blue arrowheads and neurons of the pygidium and cirri – purple arrowheads) and *Collier*, exclusively expressed in the VNC (Fig. 4 E, green square brackets). At 2 dpa, the expression patterns of both markers were globally similar between LY-411575-treated and control worms (Fig. 4D1, D2, E1, E2). However, at 5 dpa, *Elav* expression was dramatically upregulated and expanded throughout the whole mesodermal compartment of the hypertrophied pygidium in treated worms, in contrast to control animals where its expression remained restricted to few cells at the base of the pygidium and in its cirri (Fig. 4 D’1, D’2, purple arrowheads). Additionally, *Elav* expression in the anterior part of the regenerated region, including in the VNC, is reduced in treated worms. Although *Collier* expression in the VNC was maintained (Fig. 4E’2, green squared brackets), an ectopic expression domain emerged labelling differentiating neurons positioned transversely within the pygidium (Fig. 4E’2, green arrowhead).

*Synaptotagmin (Syt*), a broad marker of differentiated neurons, exhibited significantly expanded expression in LY-411575-treated worms compared to controls at 5 dpa (Fig. 4F’1, F’2). Moreover, while the cholinergic marker *VAchT* and the glutamatergic marker *VGluT* were unaffected at 2 dpa (Fig. 4G1, G2, H1, H2), *VGluT* expression was distinctly extended in the hypertrophied pygidium at 5 dpa (Fig. 4H’2, purple arrowheads).

To further examine the pygidial defects in LY-411575-treated worms at 5 dpa, we combined immunolabelling for acetylated tubulin with hybridization chain reaction (HCR) for the pygidial marker *Cdx* and the marker of differentiating neurons *Elav* (Fig. 4I, J). As expected in control worms, *Cdx* is expressed in the whole ectoderm of the pygidium (Fig. 4I1, I1’’) and is absent in the anterior tissue produced by the GZ (Fig. 4I1’). *Elav* is expressed both in the VNC (Fig. 4I2, I2’) and in the ectoderm of the pygidium, as well as in the anal cirri (Fig. 4I2, I2’’). As described earlier, acetylated tubulin labelling delineates the VNC (Fig. 4I3, I3’ – white bracket), the pygidial nerve (Fig. 4I3, I3’’ – white arrowheads), the nerve net around the gut (Fig. 4I3 – white stippled lines) and the nerve extensions in the anal cirri (Fig. 4I3 – white double arrowheads). Virtual transverse sections revealed that the VNC is overlaid by *Elav+* cells in the anterior regenerating region (Fig. 4I4, I4’), while *Elav+* and *Cdx+* cells in the pygidial ectoderm enclose the pygidial nerve (Fig. 4I4’’). In LY-411575-treated worms (Fig. 4J), *Cdx* signal is abnormally found in the whole pygidium, including internal tissues (Fig. 4J1, J1’’) while *Elav* is only maintained in few cells of the thickened VNC (Fig. 4J2, J2’) but also strongly expressed in many internal cells within the hypertrophied pygidium (Fig. 4J2’’, J4’’). Those *Cdx+* and/or *Elav+* cells are encompassing the abnormal nerve projections observed in the modified pygidium (Fig. 4J4, J4’’).

Together, these findings demonstrate that Notch signaling acts as a key regulator of pygidial neurogenesis by controlling neural progenitor specification. When Notch is inhibited, neural progenitor genes are misregulated, leading to an overproduction of neurons, including glutamatergic subtypes, and ultimately resulting in a hypertrophied, disorganized pygidium.

#### Notch regulates pygidial neurogenesis through the ligand *Delta* and several *Hes* target genes

To elucidate how the Notch pathway components are involved in pygidial neurogenesis during regeneration, we assessed their expressions upon LY-411575 inhibition (Fig. 5 and Supp. Table 5). While *Notch* receptor and its ligands *DSL1* and *3* do not appear affected by the treatment (Fig. 5A, C, D), the expression of the ligand *Delta* extends throughout the hypertrophied pygidium at 2 dpa (Fig. 5B1, B2) and displays an intense ectopic expression in two lateral patches of ectodermal cells at 5 dpa (Fig. 5B’1, B’2, white arrowheads). Among the 7 *Hes*-related genes found in neurogenic structures, 4 of them show a modified expression pattern (*Hes1, 4, 6, 13,* Fig. 5E, G, H, I), while *Hes3, 10* and *Hey* are not altered in LY-411575-treated worms (Fig. 5F, I, K, J). *Hes1+* and *Hes4+* territories are markedly reduced at 2 and 5 dpa (Fig. 5E, G) – supporting the downregulation of these genes found in the differential bulk RNA-seq data at early stages (Supp. Table 5). *Hes6* expression at 5 dpa is also extremely altered in comparison to the DMSO control (Fig. 5H), with much less *Hes6+* cells in the neurectoderm plus a shift of expression in two lateral ectodermal patches (Fig. 5H’2, white arrowheads), as observed for *Delta* (Fig. 5B’2) and a central ectodermal region of the pygidium and the anus. Finally, the overexpression of *Hes13* already revealed by the RNA-seq data is supported by its extended expression at both 2 and 5 dpa (Fig. 5J2, J’2, Supp. Table 5).

**Figure 5:**
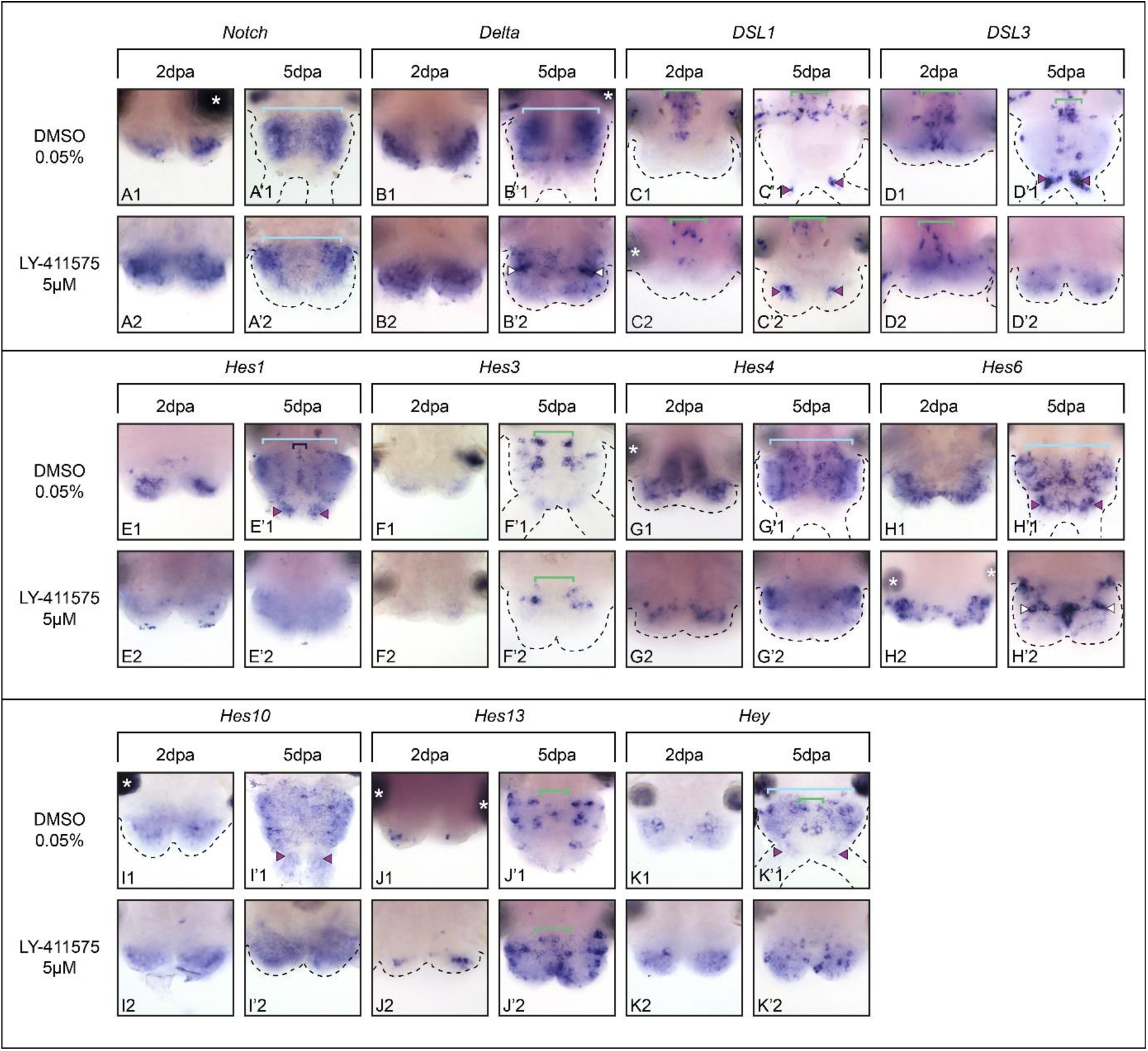
Notch inhibition alters the expression patterns of several core members of the pathway and their putative targets in the nervous system during posterior regeneration. Whole-mount *in situ* hybridizations for core components of Notch (A - D) as well as putative targets of the pathway (E-K) for LY-411575 treated worms and controls at 2 and 5dpa. Ventral views. White arrowheads point to lateral patches of ectodermal cells harboring intense expression of *Delta* and *Hes6*. Green brackets = ventral nerve cord; light blue brackets = neurectoderm; purple arrowheads = neurons of the pygidium and cirri; dark blue brackets = midline; white asterisks = non-specific staining from glands. dpa = day(s) post-amputation.

As expected, Notch inhibition also leads to significant alterations in the expression of Notch pathway components and putative *Hes* target genes expressed in non-neurogenic territories, particularly at 5 dpa (Supp. Fig. 6, Supp. Table 5). Reduced number of *Hes5+* and *Hes8+* cells are observed in LY-411575-treated worms (Supp. Fig. 6 D, E) while *Nrarp, Hes2* and *Hes8* are no longer expressed in the GZ (Supp. Fig. 6 B, C, E – yellow arrowhead).

Altogether, we demonstrated that the Notch pathway, through the action of its ligand *Delta* and several *Hes* genes, regulates the specification of neural progenitors and their differentiation into neurons in the regenerated pygidium leading to its hypertrophy upon the inhibition of the pathway. Since Notch is crucial for regenerating a functional growth zone, its inhibition impairs new tissue production and leads to an improperly formed VNC.

### Notch signaling controls central nervous system neurogenesis during post-regenerative posterior elongation in *Platynereis*

We next investigated the effects of Notch inhibition on the VNC reformation during post-regenerative posterior elongation. To circumvent the fact that Notch inhibition induces a dysfunctional regenerated GZ —characterized by minimal tissue production— we initiated LY-411575 treatments after 3 dpa, once the GZ has already reformed [29]. This staggered treatment regime (Supp. Fig. 7A) enables the formation of elongated tissues in which Notch-dependent effects on the CNS neurogenesis can be studied (Supp. Fig. 7B). Also, posterior elongation uniquely recapitulates the temporal progression of CNS and PNS neurogenesis in a postero-anterior manner: early neurogenic events occur in newly produced, growth zone–derived tissues, while later stages are visible more anteriorly.

Notch inhibition performed during posterior elongation (from 3 to 10 dpa) led to dramatic CNS defects. Neural progenitor’ markers were disrupted: *Acs2* expression is lost from superficial cells of the VNC, (Fig. 6B1, B2) while *Acs1* and *Ngn* were broadly expanded throughout the neurectoderm, escaping their normal confinement to neurogenic columns (Fig. 6A2, C2, light blue square bracket). Similarly, *Elav* and *Syt* showed expanded neuronal domains (fully differentiated or not, Fig. 6E2, F2). However, the patterns of *Collier, VAchT and VGluT* remained relatively unaffected (Fig. 6D, G, H). Acetylated tubulin labelling revealed abnormal nerve projections in the modified pygidium (Fig. 6I2, I2’’ - double white arrowheads), as previously observed at early stages. The VNC exhibited reduced neurite density, an abnormal U-shape, and was mispositioned deeper within the ventral tissues (Fig. 6I2, I2’ - white bracket). Additionally, nerve projections within the neurectoderm were highly disorganized (Fig. 6I2, I2’ - white bracket), coinciding with increased apoptosis (Supp. Fig. S8).

**Figure 6:**
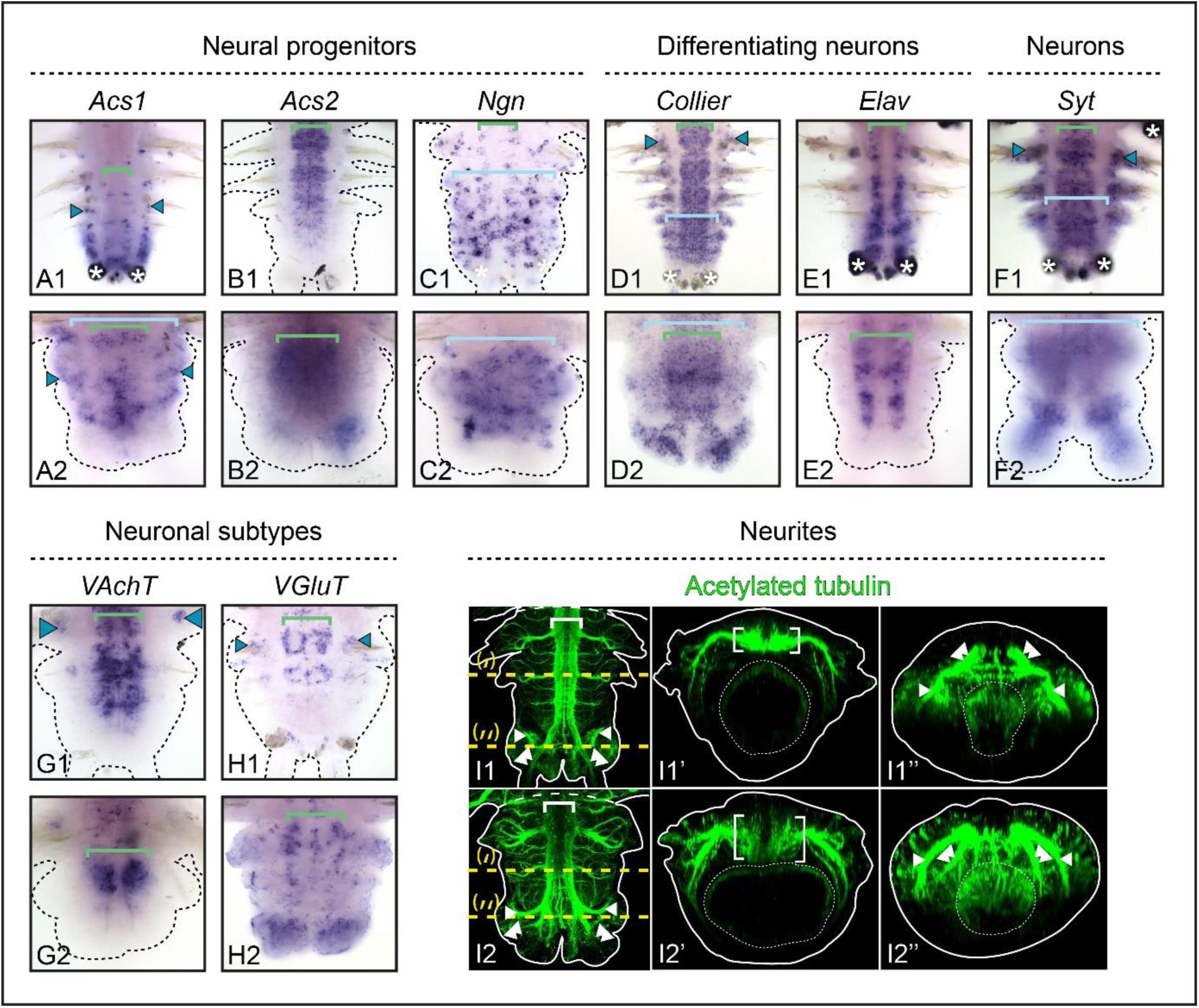
Notch pathway inhibition during post-regeneration posterior elongation leads to major neural defects in the ventral nerve chord. A - H) Whole-mount *in situ* hybridizations for markers of neural progenitors (A, -C), differentiating neurons (D, E), differentiated neurons (F) and neuronal subtypes (G, H) for LY-411575 treated worms from 3dpa to 10dpa and DMSO controls. Ventral views. Green brackets = ventral nerve cord; light blue brackets = neurectoderm; white asterisks = non-specific staining from glands. I) Acetylated tubulin immunolabelling on whole-mount regenerated parts of LY-411575-treated from 3 dpa to 10 dpa worms and controls at 10 dpa. Ventral views are on the left and corresponding virtual transverse sections along the yellow dotted lines (‘ and ‘’, respectively) are on the right. Solid white lines delineate the outlines of the samples, white dashed lines correspond to the amputation planes and white dotted lines delineate the gut. White brackets = ventral nerve cord; white arrowheads = circular pygidial nerve; white double arrowheads = cirri nerves. dpa = day(s) post-amputation.

To further assess how Notch inhibition alters the three-dimensional architecture of the neurectoderm during CNS neurogenesis, we performed HCR for key neurogenic markers (*Ngn, Elav, Syt*) known to specify larval trunk neurectoderm cells layers (Fig. 7, Supp. Fig. 9) [43]. Indeed, these genes recapitulate neurogenic events throughout the stratification of the larval neurectoderm. Superficially is a highly proliferative *Ngn+* expressing layer of neural progenitors, next is a reduced proliferative *Elav+* expressing layer of differentiating progenitors, with below a post-mitotic *Elav+* and *Syt+* expressing layer of maturing neurons and finally a VNC composed of mature *Syt+* neurons from which acetylated tubulin-labelled marked axons are projected [43]. During post-regenerative posterior elongation, neurogenesis broadly follows the same sequence of events (Fig. 7A, Supp. Fig. 9A). In control worms, the fully differentiated nuclei-dense neurectoderm (tissues far from the GZ, Fig. 7 and Supp. Fig. 9 - Transverse sections (‘)), has a distinct laminar organization. The superficial cell layers of the CNS neurectoderm express *Ngn* and *Elav* (Fig. 7A2’ - A6’), the intermediate layers co-express *Ngn, Elav* and *Syt* (Fig. 7A2’-A6’; Supp. Fig. 9A2’-A5’), while the deeper ones express *Elav* and *Syt* (Supp. Fig. 9A2’-A5’). Finally, a *Syt+* cell layer corresponds to the VNC (white brackets) from which neurites are projected (Fig. 7A6’, Supp. Fig. 9A3’). Lateral nerves innervate the ganglions of the PNS, which are *Ngn+, Elav+* and *Syt+* (Fig. 7A1’-, A6’; Supp. Fig. 9A1’-A5’, blue arrowheads). During early neurogenesis (tissues close to the GZ, Fig. 7 and Supp. Fig. 9 - Transverse sections (‘’)), superficial thin layers of the CNS neurectoderm are composed of *Ngn+* and *Elav+* cells (Fig. 7A2’’-A6’’), while deeper cells are co-expressing *Ngn, Elav* and *Syt* (Fig. 7A2’’-A6’’; Supp. Fig. 9A2’’-A5’’). As for late neurogenesis, the VNC (white brackets) is positioned below the whole CNS neurectoderm, while lateral nerve extensions support the PNS neurectoderm anlagen composed of *Ngn+* and *Elav+* cells Fig. 7A1’’-A6’’; blue arrowheads). Sagittal sections (Fig. 7 and Supp. Fig. 9 – (‘’’)) illustrated that *Ngn* (Fig. 7A2’’’, A4’’’-,A6’’’) is the earliest marker to be expressed in superficial neurectodermal cells, with its expression diminishing as differentiation proceeds. Conversely, *Elav* and *Syt* are expressed subsequently and maintained throughout CNS maturation (Fig. 7A3’’’-A6’’’, Supp. Fig. 9A2’’’-A5’’’).

**Figure 7:**
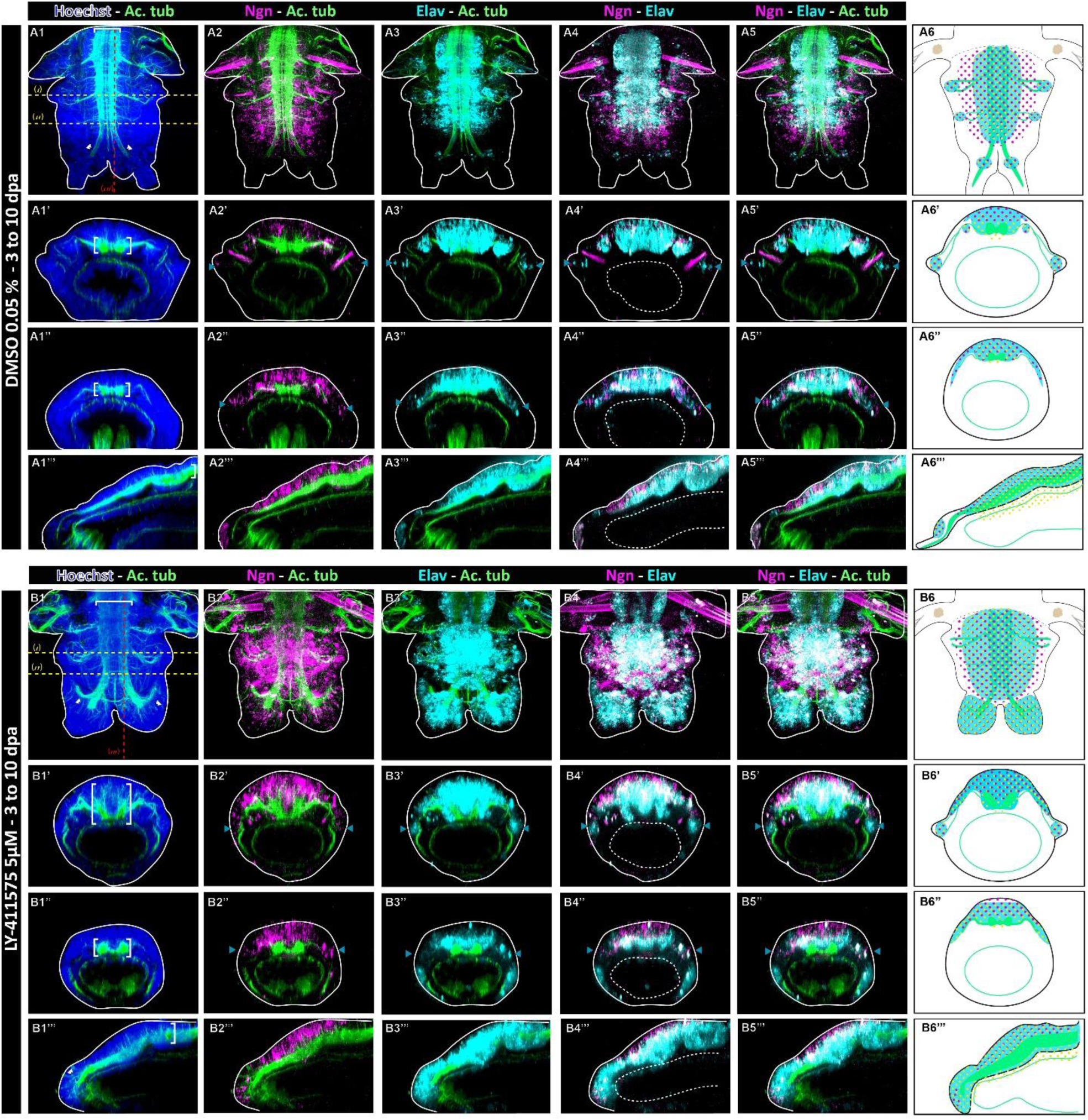
Notch pathway inhibition during post-regeneration posterior elongation leads to a disorganized thicker ventral nerve cord and to the enlargement of the neurectodermic territories. Hybridization chain reactions (HCR) for *Elav* (cyan) and *Ngn* (magenta) coupled with immunolabelling for acetylated tubulin (green) and nuclei staining with Hoechst (blue) for controls at 10 dpa (top) and LY-411575 treated regenerated parts from 3 dpa to 10 dpa (bottom). Ventral views are on top for each condition and corresponding virtual transverse sections (along (‘) and (‘’) in yellow) and sagittal section ((‘’’) in red) are at the bottom. On the right, schematic drawings depict the expression patterns of *Elav* (cyan), *Ngn* (magenta dots) and *Syt* (yellow dots – from Supp. Figure 9) as well as their arrangement around the main structures of the central nervous system and peripheral nervous system (green). Solid white lines delineate the outlines of the sample, and white dotted lines delineate the gut. White brackets = ventral nerve cord; white double arrowheads = cirri nerves; blue arrowheads = PNS. dpa = day(s) post-amputation.

In the context of Notch inhibition, HCR and acetylated-tubulin co-labelling confirmed the drastic alteration of the regenerated neural structures (Fig. 7B and Supp. Fig. 9B). In a fully differentiated neurectoderm of LY-411575-treated worms (tissues far from to the GZ, Transverse section (‘)), the abnormally U-shaped VNC (white brackets) is overlaid by a markedly thicker nuclei-dense CNS neurectoderm (Fig. 7 B1’, Supp. Fig. 9B1’), whose most of the cells are *Ngn+, Elav+* and *Syt+* (Fig. 7 B6’). This territory extends on superficial cells until joining the PNS ganglions (Fig. 7 B6’). At the level of the VNC, the deepest cell layers of this laterally and thickly enlarged structure contain only *Elav+* and *Syt+* cells (Fig. 7 B6’). During early neurogenesis (tissues close to the GZ, Transverse section (‘’)), the two cords of the VNC are slightly apart and the neurectoderm is already slightly thickened (Fig. 7 B1’’, Supp. Fig. 9B1’’, white brackets). This CNS neurectoderm is composed of cell layers that are, from top to bottom, *Ngn+*, then *Ngn+, Elav+* and *Syt+* and finally *Elav+* and *Syt+* (Fig. 7 B6’’). All of these three markers appear to be expressed almost concomitantly in very recently-produced cells from the GZ and their expression is broadly maintained throughout CNS postero-anterior differentiation (Fig. 7 B6’’’). Hence, Notch inhibition disturbs the dynamic of the neurogenic cascade at the neural progenitor determination step, leading to a thickened CNS neurectoderm and an abnormally-shaped VNC with neurite defects.

We then dissected how Notch pathway components regulate CNS neurogenesis (Fig. 8) and showed that while *Notch* receptor and the ligands *DSL1* and *3* do not appear much affected by the treatment (Fig. 8A, C, D), the ligand *Delta* expression is enhanced in the CNS neurectoderm at 10 dpa (Fig. 8B). Upon Notch-inhibition, 3 of the 7 *Hes*-related genes expressed in neurogenic structures appear not affected (*Hes3, 6* and *10,* Fig. 8f, G, H) while *Hes1* and *Hes4* are down-regulated (Fig. 8E, G). Expression of *Hes1* is maintained in the midline (Fig. 8E2 – dark blue square bracket). Interestingly, *Hes13* and *Hey* expression domains are extended in LY-411525 treated worms (Fig. 8J, K). Many more *Hes13+* cells are found in the whole ventral surface of the regenerated structure, likely in both the CNS and PNS, in a salt and paper fashion, in comparison to controls (Fig. 8J). *Hey* expression extends to the whole CNS neurectoderm in a disorganized manner when Notch is inhibited (Fig. 8K – light blue square bracket). Additional data on Notch pathway members altered expressions in LY-411525 treated worms show that Notch is not restricted to neurogenic functions during post-regenerative posterior elongation. *Delta*, *Hes2* and *Nrarp* expressions in the GZ are lost in LY-411575 treated worms (Fig. 8B2, Supp. Fig 10A, B – yellow arrowheads) supporting the idea that Notch regulates not only its regeneration but also partly its functioning. In addition, Notch determines the identity of the two main cell types composing the chaetal sacs, the follicle cells and chaetoblasts [27, 44]. While *Nrarp* and *Hes2* expressions in the follicle cells appears reduced or lost upon Notch inhibition (Supp. Fig. 10A2, B2 - pink arrowheads), *Hes12* expression in the chaetoblast is extended, and *Hes12*+ cells are even present in the pygidium (Supp. Fig. 10C2 - pink arrowheads).

**Figure 8:**
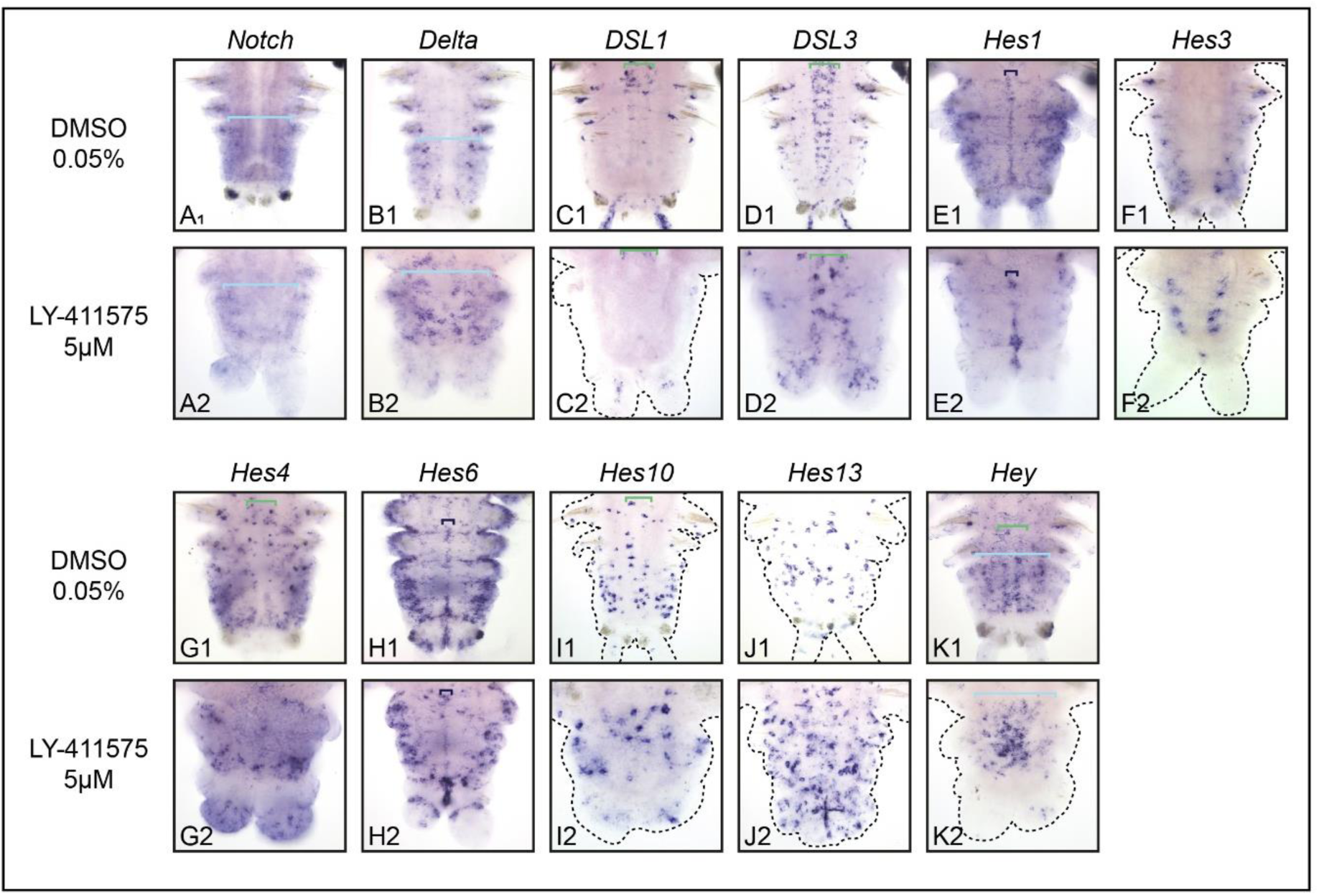
Notch inhibition during post-regeneration posterior elongation alters the expression patterns of several core members of the pathway and their putative targets in the nervous system. Whole-mount *in situ* hybridizations for core components of Notch (A to D) as well as putative targets of the pathway (E to K) for LY-411575-treated worms from 3 dpa to 10 dpa and controls at 10 dpa. Ventral views. Green brackets = ventral nerve cord; light blue brackets = neurectoderm; purple arrowheads = cirri; dark blue brackets = midline; white asterisks = non-specific staining from glands. dpa = day(s) post-amputation.

In conclusions, we definitively demonstrated that the Notch pathway regulates neurogenesis during both pygidial regeneration and CNS patterning during posterior elongation.

## DISCUSSION

Notch signaling is a key cell-cell communication pathway that orchestrates diverse cellular behaviors in both embryonic and post-embryonic developmental contexts across vertebrates and ecdysozoans. Its wide range of functions is enabled by the modular interactions among receptors, ligands, and a variety of target genes [4, 5, 45]. However, the role of Notch signaling in spiralians remains relatively unexplored, which has limited our understanding of its ancestral functions at the bilaterian scale. Here we uncovered the multiple functions of Notch signaling during regeneration and posterior elongation in the annelid *Platynereis*. This spiralian model possesses a complete set of core Notch components— including the receptor *Notch*, ligands *Delta* and *Jagged,* the regulator *Nrarp* and an array of *Hes/Hey* effectors [8, 46]. Our study demonstrates that the *Platynereis* Notch pathway is remarkably modular, participating in several distinct processes. We show that Notch signaling is essential for the proper regeneration of the growth zone stem cells, for the formation of bristles during posterior elongation (as previously described during larval development [27]) and for the regulation of neurogenesis during both post-embryonic processes likely *via* the transcriptomic modulation of up to eight *Hes/Hey* genes. In particular, our comprehensive molecular analysis of the neurogenic cascade involved in pygidial regeneration and ventral nerve cord formation through posterior elongation, revealed that impairing Notch signaling leads to excessive neurogenesis. This indicates that Notch signaling plays a pivotal role in maintaining the balance between neural progenitor specification and the production of differentiated neurons. Surprisingly, previous research on Notch signaling did not find supporting evidence for such effects during early larval neurogenesis [27].

In vertebrates and *Drosophila*, Notch signaling is well known for its function in neuronal cell fate specification, where it preserves a pool of progenitors while directing differential cell fates, such as neurons *versus* glial cells [14, 15]. In *Platynereis*, it remains unclear whether the excess neurons produced upon Notch signaling inhibition comes at the expense of other cell types (*e.g.* glial cells), due in part to the current lack of a specific glial marker. Nevertheless, our data support a conserved and ancestral role for Notch signaling in regulating neuronal balance across bilaterians. Interestingly, a recent study in the planarian *Schmidtea* revealed that Notch signaling is also regulating glial specification during regeneration via the interactions between mature neurons and non-neural progenitors, suggesting a conserved role of Notch signaling in glial development specification rather than on neuronal balance [47]. Discriminating between these two hypotheses will require more research in additional spiralian models.

Our study also illustrates an essential role of Notch signaling in regulating axon guidance and neurite growth. *Platynereis* worms have a complex VNC and pygidial neural network composed of highly organized neuropils that rely on precise axon guidance mechanisms [43]. In the absence of Notch signaling, we observed aberrant axonal projections during both pygidial nerve regeneration and VNC formation in posterior elongated tissues, leading to a scattered and enlarged VNC. We found a concomitant increase of cell deaths by apoptosis, possibly due to the failure of misrouted axons to establish proper target connections [48]. Although Notch has been shown to regulate axon patterning and neurite growth in both vertebrates [19, 20] and *Drosophila* [16–18]. These findings suggest that Notch-mediated axon guidance might represent an ancestral function in protostomes and even bilaterians.

Our work highlights the multifaceted and conserved roles of Notch signaling in bilaterian neurogenesis, ranging from the upstream neural progenitors’ specification to the ultimate axon guidance step allowing fine neural circuitry. These findings underscore the need for further studies in diverse model organisms to fully elucidate its evolutionary and developmental significance.

## MATERIALS AND METHODS

### *Platynereis dumerilii’s* culture, amputation procedure and biological material fixation

*Platynereis* juvenile worms were obtained from a husbandry established at the Institut Jacques Monod (for detailed breeding conditions see [49, 50]). Standard worms used in experiments were 3-4-month-old with 30-40 segments and were amputated according to the procedure described previously [29, 50]. For the majority of experiments performed (*i.e. in situ* hybridizations, antibody stainings, EdU and TUNEL assays as well as hybridization chain reactions (HCR) – see below for each detailed procedure), regenerative parts at the desired stage and condition =were collected and fixed in 4% paraformaldehyde (PFA) diluted in PBS Tween20 0.1% (PBT) for 2 hours at room temperature (RT). Following fixation, whole mount samples were washed in PBT, gradually transferred in 100% methanol (MeOH) then stored at −20°C [50]. For phalloidin staining (see below), after fixation without MeOH dehydration, regenerative parts were stored in PBT at 4°C for up to 4 days prior to labelling [29].

### Histologic samples fixation and sectioning

Histological sections were performed as described in [51]. Briefly, samples were fixed in 4% PFA diluted in PBS 1X for 1h30 at RT, washed in PBS 1X, cryoprotected in a solution of PBS/sucrose 30% for 4-5 days at 4°C and then transferred into OCT embedding medium (Tissue Freezing Medium, Leica). Next, samples were put into molds and positioned according to the desired type of section (longitudinal), then frozen with dry ice and stored at −80°C. Samples were cut using a microtome (Leica CM3050S) and sections of 12-14µM were collected on SuperFrost glass slides prior to storage at −80°C.

### Whole-mount *in situ* hybridization (WMISH), antibody staining and phalloidin labelling

Colorimetric NBT/BCIP WMISH and immunolabelling were performed as previously described [43, 50]. For all experiments, following rehydration, samples were treated with 40 µg/ml proteinase K in PBT for 10min, 2mg/ml glycine PBT for 1min, 4% PFA PBT for 20min and washed in PBT prior to hybridization or labelling. Neurites labelling was done as previously described [43], using the mouse anti-acetylated tubulin (Sigma 1:500) antibodies and fluorescent secondary anti-mouse IgG Alexa Fluor 488 or 555 conjugate (Invitrogen, 1:500). For phalloidin labelling, samples were incubated in phalloidin-Alexa 555 (Molecular Probes, 1:100) overnight at 4°C. Next, samples were nuclei counter-stained with Hoechst 0.1% overnight at 4°C and mounted in glycerol/DABCO (2.5mg/ml DABCO in glycerol) for confocal imaging (see below).

### EdU cell proliferation and TUNEL cell death assays

Proliferating cells were labelled by incubating worms with 5 µM of the thymidine analog 5-ethynyl-2′-deoxyuridine (EdU), for 1h in natural fresh sea water prior to fixation. Various incubation conditions (duration and biological stage) and pulse and chase experiments were performed as described in the Results section and related figures. Fixed samples were subsequently fluorescently labelled with the Click-it EdU Imaging Kit (488 or 555 nm, ThermoFisher) as previously described in [50]. TUNEL labelling was performed using the Click-iT TUNEL kit (647 nm, ThermoFisher), as previously described in [38]. In both cases, samples were nuclei counter-stained with Hoechst 0.1% overnight at 4°C and mounted in glycerol/DABCO (2.5mg/ml DABCO in glycerol) for confocal imaging (see below).

### Hybridization Chain reactions (HCR)

HCR coupled with immunolabelling was implemented in *Platynereis* following the HCR 3.0 protocol developed in [52]. More specifically, the primary antibody incubation of the immunolabelling was performed simultaneously with the amplification step of HCR. Up to 25 couples of probes were designed using an in-house ProbeMaker based on the HCR 3.0 Probe Maker v1.0 (http://hcr.wustl.edu/input/quick/) [53]. Each probe sequence was then manually blasted against *Platynereis’*s transcriptome, and probes matching several genes were removed. All validated probes were combined in an oligo pool ordered at IDT (see probes list in Supplementary Table 6). Given the variety of adapters (Molecular Instruments) that can be used with different fluorophores, we made the following selection: *Cdx, Ngn* and *Syt* with the adapter B1 fused with Alexa 647, and *Elav* with the adapter B3 fused with Alexa 594.

### Images acquisition, treatments and analyses

Bright-field images of colorimetric WMISH samples were acquired with a Leica CTR 5000 microscope. Fluorescent confocal images of samples/ sections were acquired with either a Zeiss LSM780 or LSM980 confocal microscopes. Image processing (contrast and brightness, z-projection, auto-blend layers, transversal and sagittal views) was performed using FIJI and Abode Photoshop. Figures were assembled with Adobe Illustrator. EdU and TUNEL cell counts were performed using IMARIS 9.5.0 (Oxford Instruments) following the automatic cell counting procedure defined in [38, 51].

### Notch signalling pathway inhibitors treatments, scoring and statistical analyses

Chemical inhibitions of the Notch signalling pathway were performed using three different inhibitors, widely used to specifically block the gamma-secretase complex responsible for the cleavage of Notch, thus preventing transcription of the target genes: DAPT, LY-411575 and RO-4929097 [54]. These inhibitors have been previously successfully used to disrupt Notch signaling in *Platynereis* embryos and larvae [27], but also in a diversity of organisms during both development and regeneration [55–61]. We first determined the efficient concentrations for each inhibitor by performing treatments with different concentrations (1, 5 and 10 µM for LY-411575, 5 and 10 µM for RO-4929097 and 40 µM for DAPT, based on [27], from a 10mM stock solutions in DMSO and in comparison to DMSO controls). Upon amputation, we immediately incubated the worms in 2 ml of each inhibitor solution on 12-well plate and assessed the effects by scoring the regenerative stages reached by each worm every day for 5 days of treatment (when posterior regeneration is over) [29, 50]. Control worms were incubated in natural fresh sea water with identical concentrations of DMSO. All solutions were refreshed every 24 hours to maintain their activities for the whole duration of the experiments [50]. Thus, 5µM for both LY-411575 and RO-4929097 and 40 µM for DAPT, were found to be the most effective concentrations, resulting in a consistent and reproducible effect: blocking regeneration around stage 2 (Supp. Fig. 2A, B and C respectively). As all inhibitors led to similar effects on regeneration, we decided to pursue our experiments using only LY-411575 at 5µM.

All statistical tests and subsequent graphical representations were performed using GraphPad Prism 9. Mann–Whitney U tests were used to compare samples between different experiments/conditions. The following symbols are used to indicate statistical significance in figures: ns p>0.05; * p≤0.05; ** p≤0.01; *** p<0.001; **** p≤ 0.0001).

#### Determination of gene expression levels during posterior regeneration

In an updated version of our *Platynereis* reference transcriptome [35] (Supp. Table 2), 26 genes corresponding to both the Notch signaling pathway core machinery and their putative target genes from the Hairy enhancer of split multigenic family [8, 27] were identified. Their respective expression levels were determined during the course of regeneration (*i.e.* stages 0, 1, 2, 3, 5 days post amputation as well as non-amputated control – 2 to 3 replicates) using our previously produced RNA-seq datasets of posterior regeneration [35] (Supp. Table 1). Expression level dynamics were visualized using heatmap.2 from the gplots R package.

#### Transcriptomic analysis of Notch inhibition effects on posterior regeneration

*Sample production and collection*: 1 dpa and 2 dpa samples treated with 5 µM of LY-411575 and 0.05% DMSO controls were produced. For each stage and condition, three biological replicates per stage and condition were produced independently, each one containing 200 regenerating parts recovered with as little non-amputated tissue as possible (typically half a segment).

*RNA extraction, library construction and sequencing*: For all samples (n=12), total RNA was extracted and their quality was assessed as described previously [35]. Libraries and Illumina TruSeq Stranded sequencing (75 bp in single-end) were performed at the Ecole normale supérieure GenomiqueENS core facility (Paris, France), as detailed in [35]. All raw reads from individual sequencing libraries are deposited in the European Nucleotide Archine (ENA) (Supp Table 2).

*Read processing and mapping:* Reads were quality checked using FastQC v0.11.8 and trimmed for low quality reads and adapter using fastp v0.23.2. Kallisto v0.48.0 within the Trinity v2.13.2 toolkit [62] was then used to perform pseudo-mapping and quantification of the reads on the updated reference transcriptome and to generate the raw count matrix. The raw count matrix was processed using EdgeR v3.40.2 [63] to obtain a TMM count matrix. The intersection of expressed genes between each condition was plotted using the UpSetR v1.4.0 R package [64]. PCA plots were performed based on the processed raw count matrix to a count per million matrix. Only genes with a TMM value superior or equal to 1 TMM were retained. The count per million matrix was batch-corrected using the removeBatchEffect function from EdgeR. The PCA plot was performed using the factoextra R package v1.0.7.

*Differentially expressed genes identification*: For each treated *versus* control condition (at 1 and 2 dpa), differential expression analyses were conducted using the EdgeR R package based on the raw count matrix. Only genes with a TMM value superior or equal to 1 TMM in at least one condition were considered. A design matrix was built to perform batch correction. The p-values were adjusted for multiple tests using the Benjamini-Hochberg FDR correction. Genes with a adjusted p-value less than 0.05 and an absolute log fold change value of at least 1 were considered as differentially expressed genes (DEGs). Volcano plots were constructed using the EnhancedVolcano v1.16.0 R package [65]. DEGs were annotated with Trinotate [66] and from the top homology blast on the mouse proteome, as previously described [35]. We used the VennDiagram package in R [64] to quantify and visualize shared DEGs between comparisons.

*Gene ontology and enrichment analysis for differentially expressed genes:* We performed Gene Ontology (GO)-term enrichment analyses on DEG lists using clusterProfiler [67]. For the DEGs, a full list of enriched GO terms is provided in Supp Tables 3 and 4 and the top 20 per comparison for the category “Biological processes” are presented on dotplots.

## Supplementary material

**Supplementary Figure 1:**
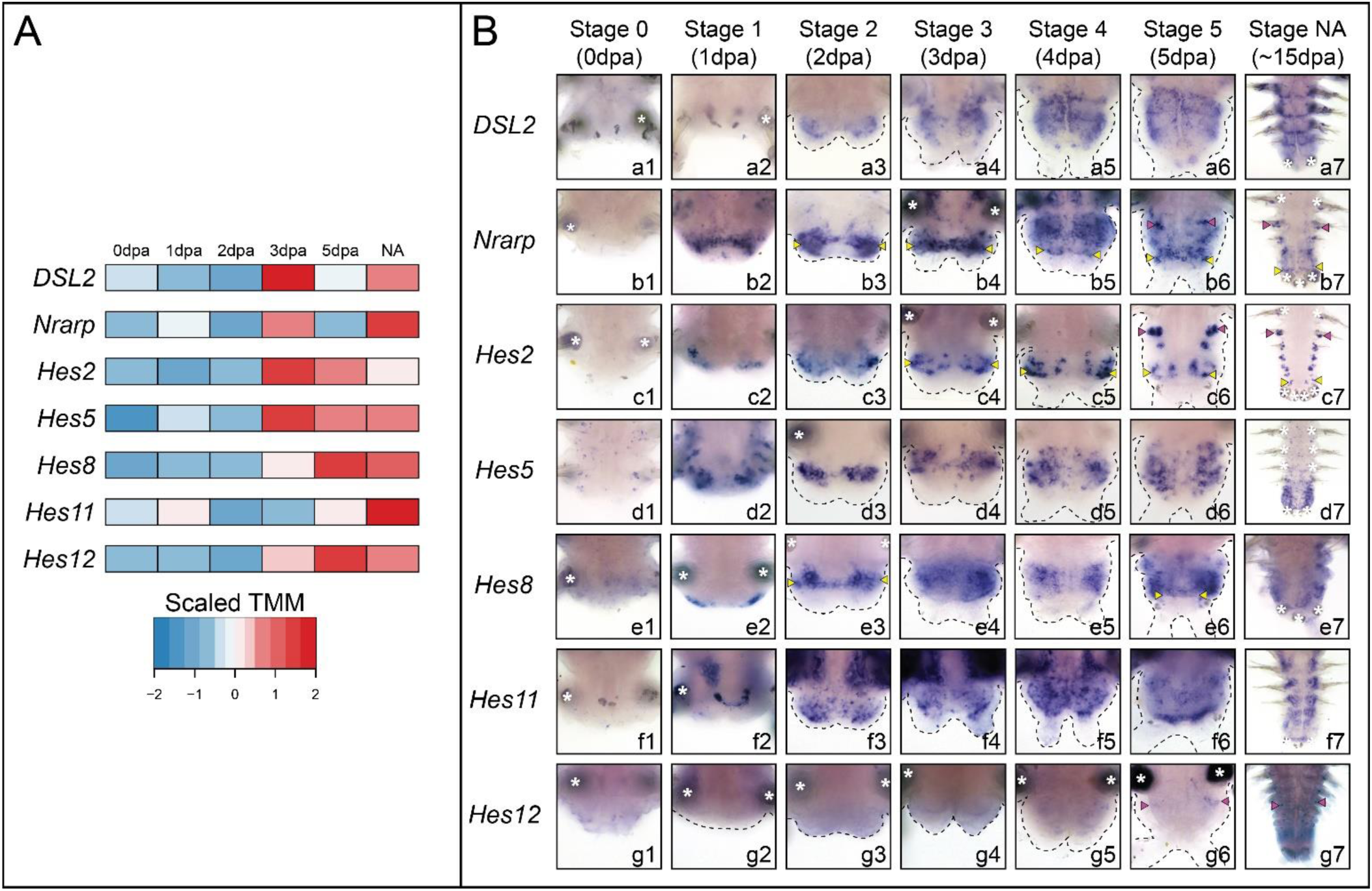
Dynamic expression of core members of the Notch pathway and its putative target genes in non-neurogenic territories during posterior regeneration. A) Heatmap representation of expression levels of several Notch components and *Hes* genes during posterior regeneration [35]. B) Whole-mount *in situ* hybridizations (ventral views) of Notch components and *Hes* genes expressed in non-neurogenic structures during regeneration. Yellow arrowheads = growth zone involved in posterior elongation of the animals [28]; pink arrowheads = chaetal sacs producing the parapodial bristles; white asterisks = non-specific staining from glands. dpa = day(s) post-amputation, NA = non-amputated.

**Supplementary Figure 2:**
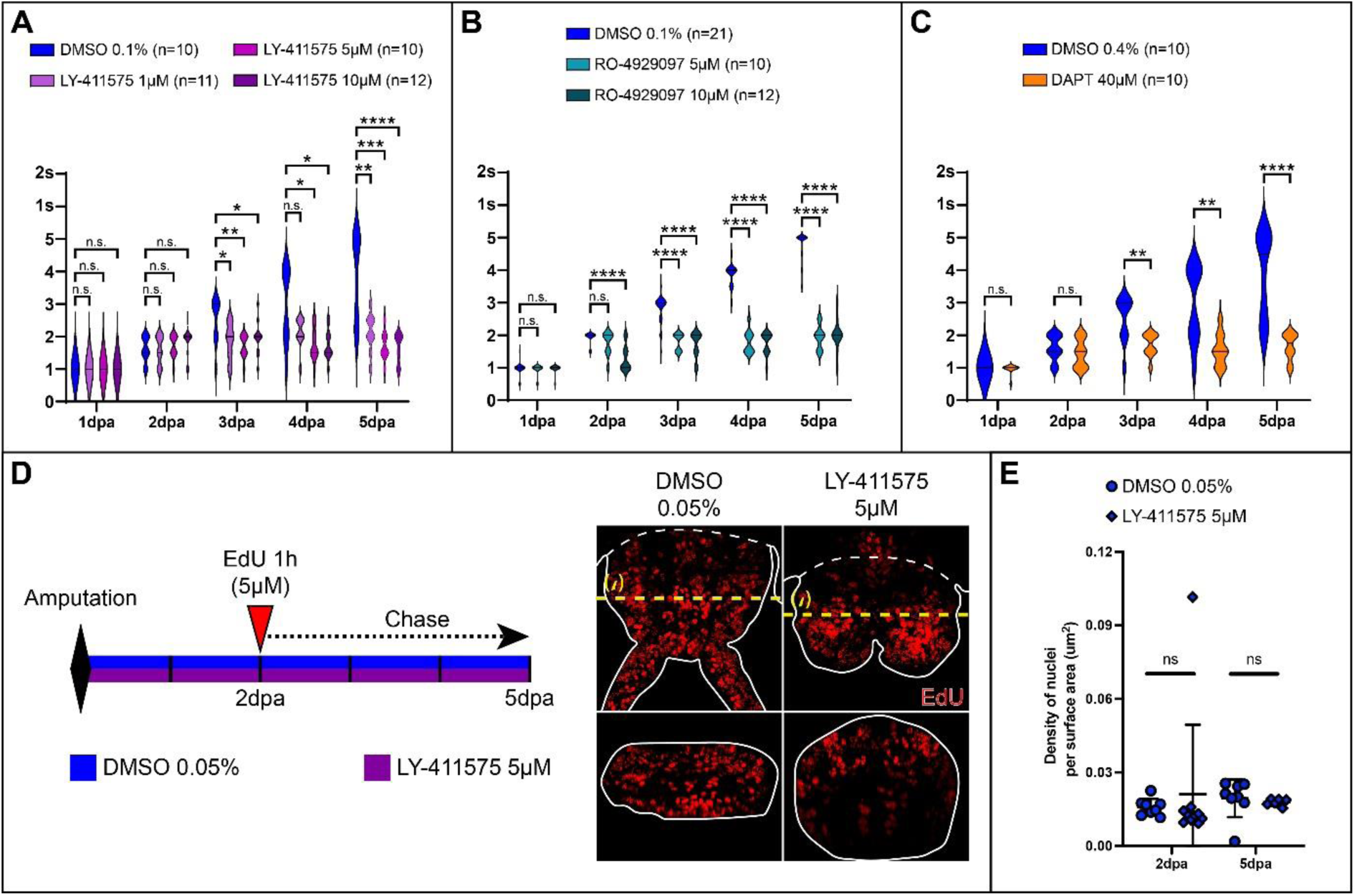
Determination of the optimal Notch inhibitors concentrations and additional morphological and cellular effects of LY-411575 treatment during posterior regeneration in the annelid *Platynereis*. A - C) Violin plots representing the regeneration stages reached by each worm every day for 5 days of treatments. A) LY-411575 treatments at 1, 5 and 10 µM in comparison to control (DMSO 0.1%); B) RO-4929097 treatments at 5 and 10 µM in comparison to control (DMSO 0.1%); C) DAPT treatment at 40 µM in comparison to control (DMSO 0.4%). n = number of worms used per experiment. D) Schematic representation of the EdU chase experiment during Notch inhibition and corresponding EdU labelling on LY-411575-treated worms and controls. Ventral views are on top and corresponding virtual transverse sections (along the yellow dotted lines) are at the bottom. E) Comparison of density of nuclei per surface area for LY-411575-treated worms and controls at 2 and 5 dpa. Solid white lines delineate the outlines of the samples and white dashed lines correspond to the amputation planes. Each bracket corresponds to a Mann-Whitney U test. n.s. p >0.05; * p<0.05; ** p<0.01; *** p<0.001; **** p<0.0001. dpa = day(s) post-amputation.

**Supplementary Figure 3:**
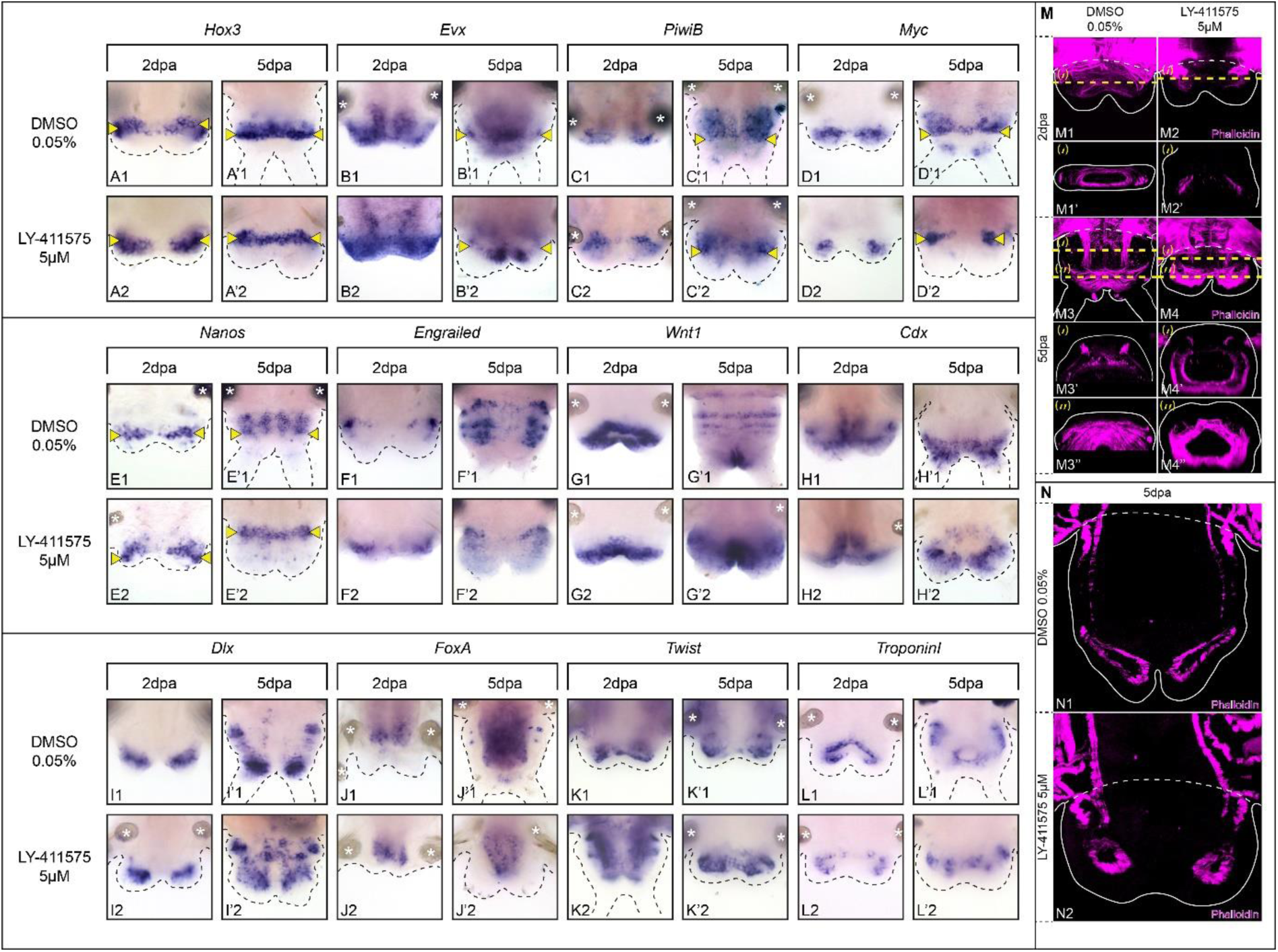
Effects of Notch signaling pathway inhibition on several tissues during posterior regeneration in *Platynereis*. A-L) Whole-mount *in situ* hybridizations for markers of the growth zone (A, B), stem cells (C-E), segmentation (F, G), pygidium (H), pygidial cirri and appendages (I), gut (J) and muscles (K, L) for LY-411575 treated worms and controls at 2 and 5 dpa. Ventral views. M) Phalloidin labelling on whole-mount regenerated parts of LY-411575-treated worms and DMSO controls at 2 and 5 dpa. Ventral views are on top and corresponding virtual transverse sections (along the yellow dotted lines are at the bottom. N) Phalloidin labelling on longitudinal cross-sections of LY-411575-treated worms and controls at 5dpa. M and N) Solid white lines delineate the outlines of the samples, white dashed lines correspond to the amputation planes. Yellow arrowheads = growth zone involved in posterior elongation of the animals [28]; white asterisks = non-specific staining from parapodial glands. dpa = day(s) post-amputation.

**Supplementary Figure 4:**
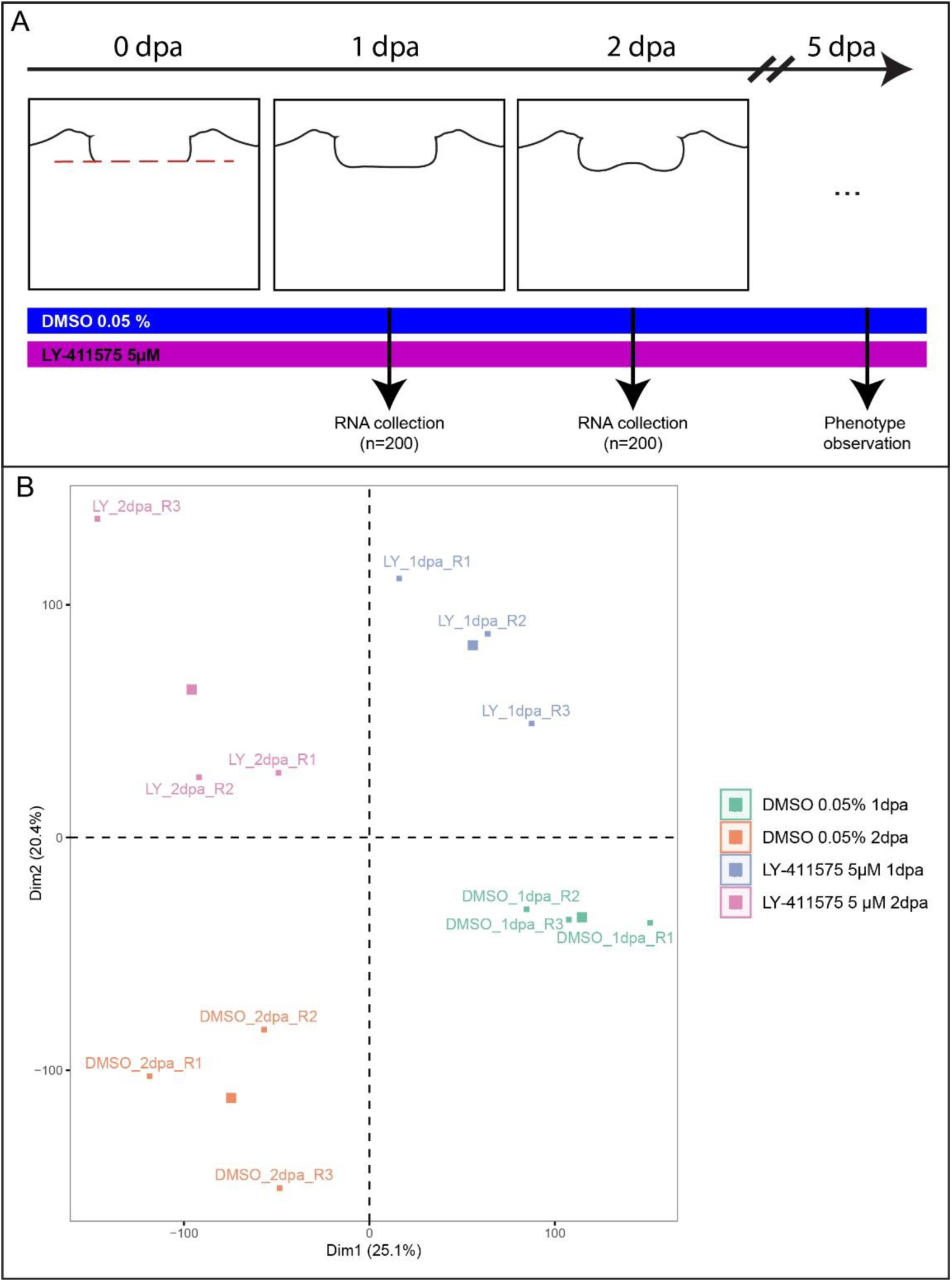
Schematic representation of the RNA-seq experiment. A) RNA-seq experiment design. Total mRNA was extracted from regenerative parts of LY-411575-treated worms and DMSO controls at 1 and 2 dpa, when the structures are still morphologically comparable. Three replicates were produced and about 200 regenerating parts (plus half a segment) were collected per replicate. Phenotype was observed at 5 dpa for a couple of animals to ensure the quality of each batch. B) PCA analysis of the 12 RNA-seq samples (4 conditions with 3 replicates). dpa = day(s) post-amputation.

**Supplementary Figure 5:**
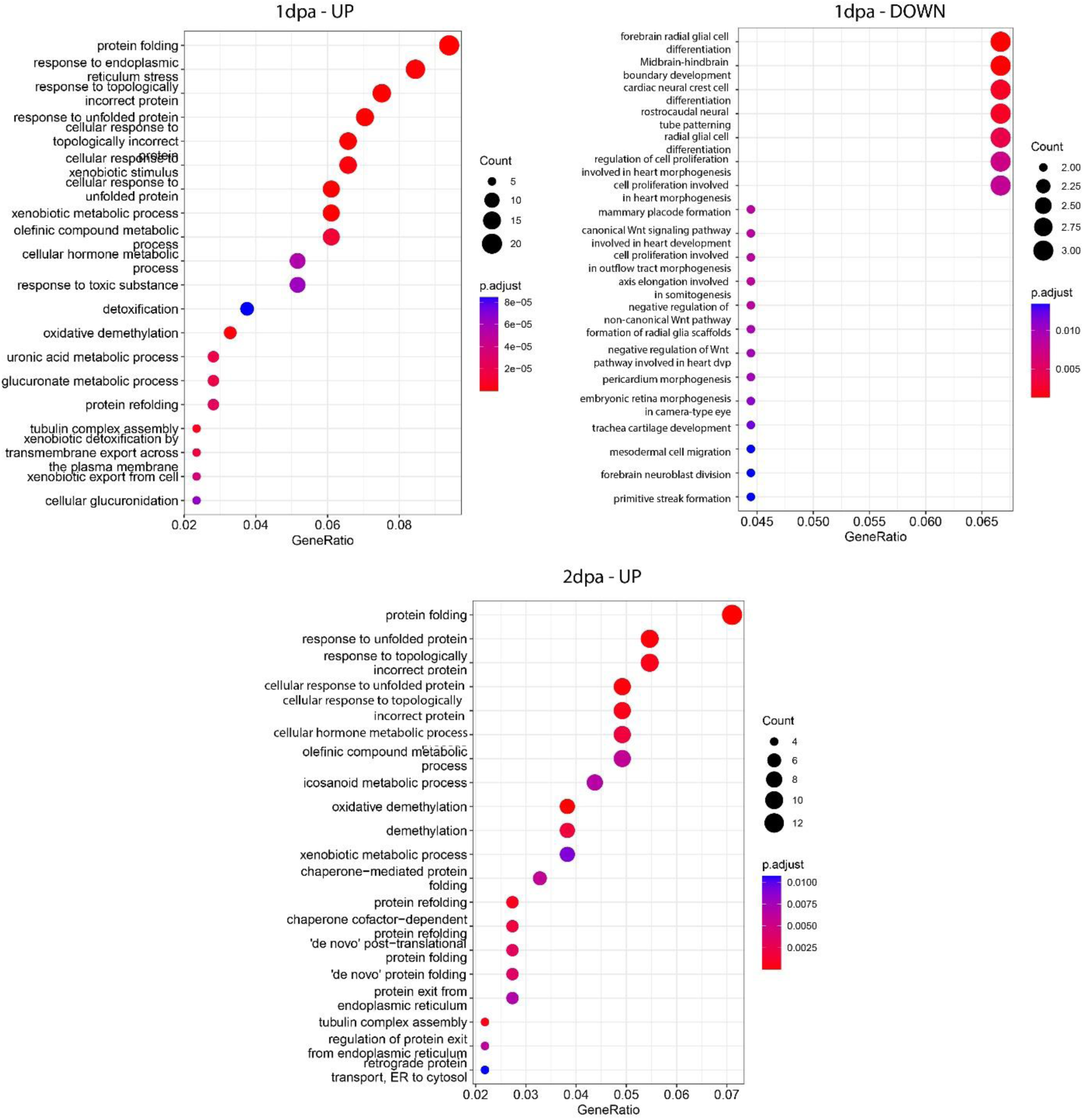
Gene ontology enrichment analysis of differentially expressed genes (DEGs) between LY-411575-treated animals and controls. Dotplots showing most significant over-represented Biological Process GO terms (Top 20) for DEG upregulated at 1 dpa in LY-treated conditions (top left), downregulated at 1 dpa (top right) and upregulated at 2 dpa (bottom). Circles area is proportional to the fraction of transcripts in each condition falling into the corresponding GO term (lines), colors correspond to the adjusted p-value of the enrichment. Gene ratio in x-axis corresponds to the percentage of genes found in each GO term over the total number of genes for each condition.

**Supplementary Figure 6:**
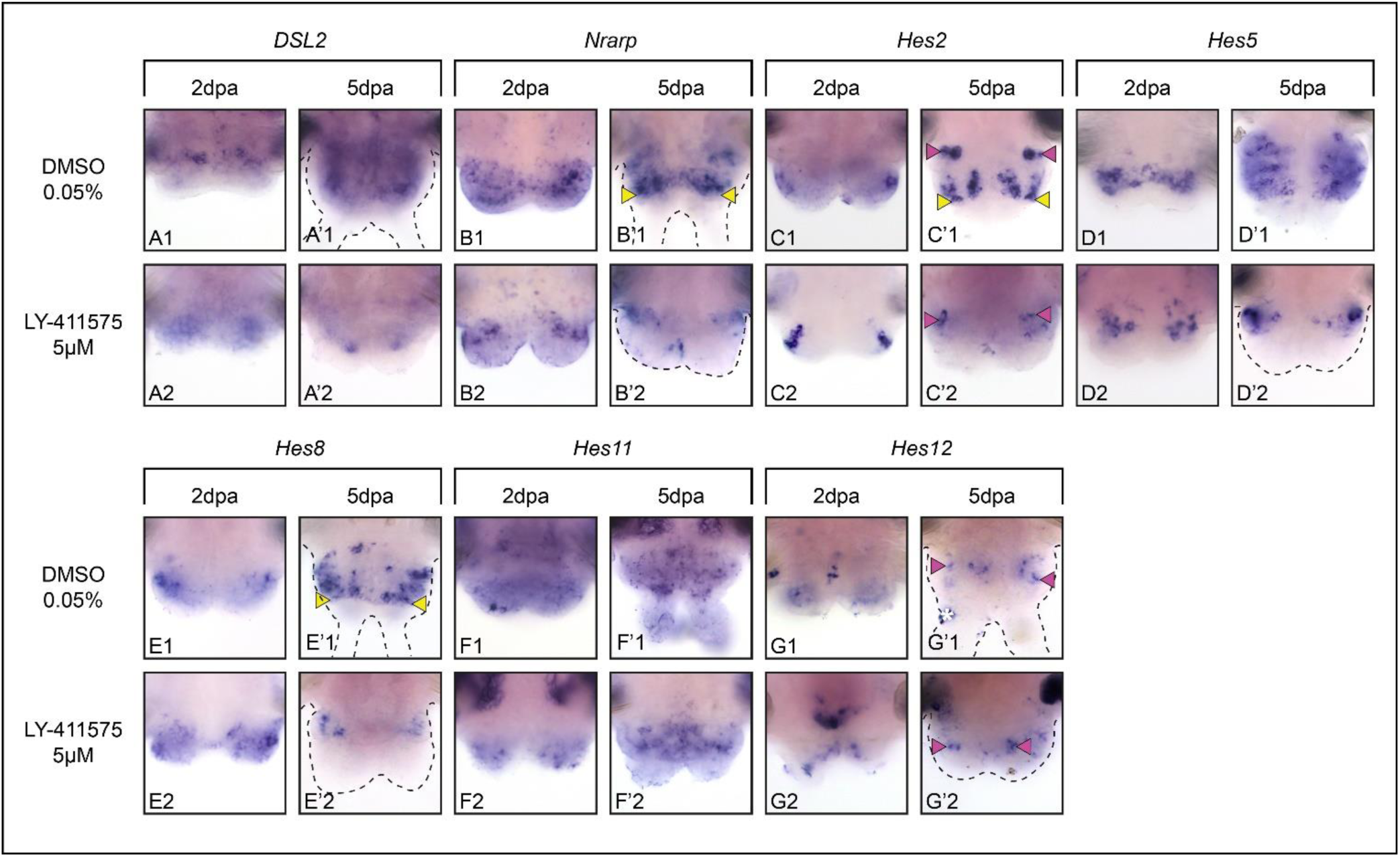
Impact of Notch inhibition on non-neural territories during posterior regeneration. Whole-mount *in situ* hybridizations of Notch components and *Hes* genes expressed in non-neurogenic structures for LY-411575 treated worms and DMSO controls at 2 and 5 dpa. Ventral views. Yellow arrowheads = growth zone involved in posterior elongation of the animals [28]; pink arrowheads = chaetal sacs producing the parapodial bristles; white asterisks = non-specific staining from glands. dpa = day(s) post-amputation.

**Supplementary Figure 7:**
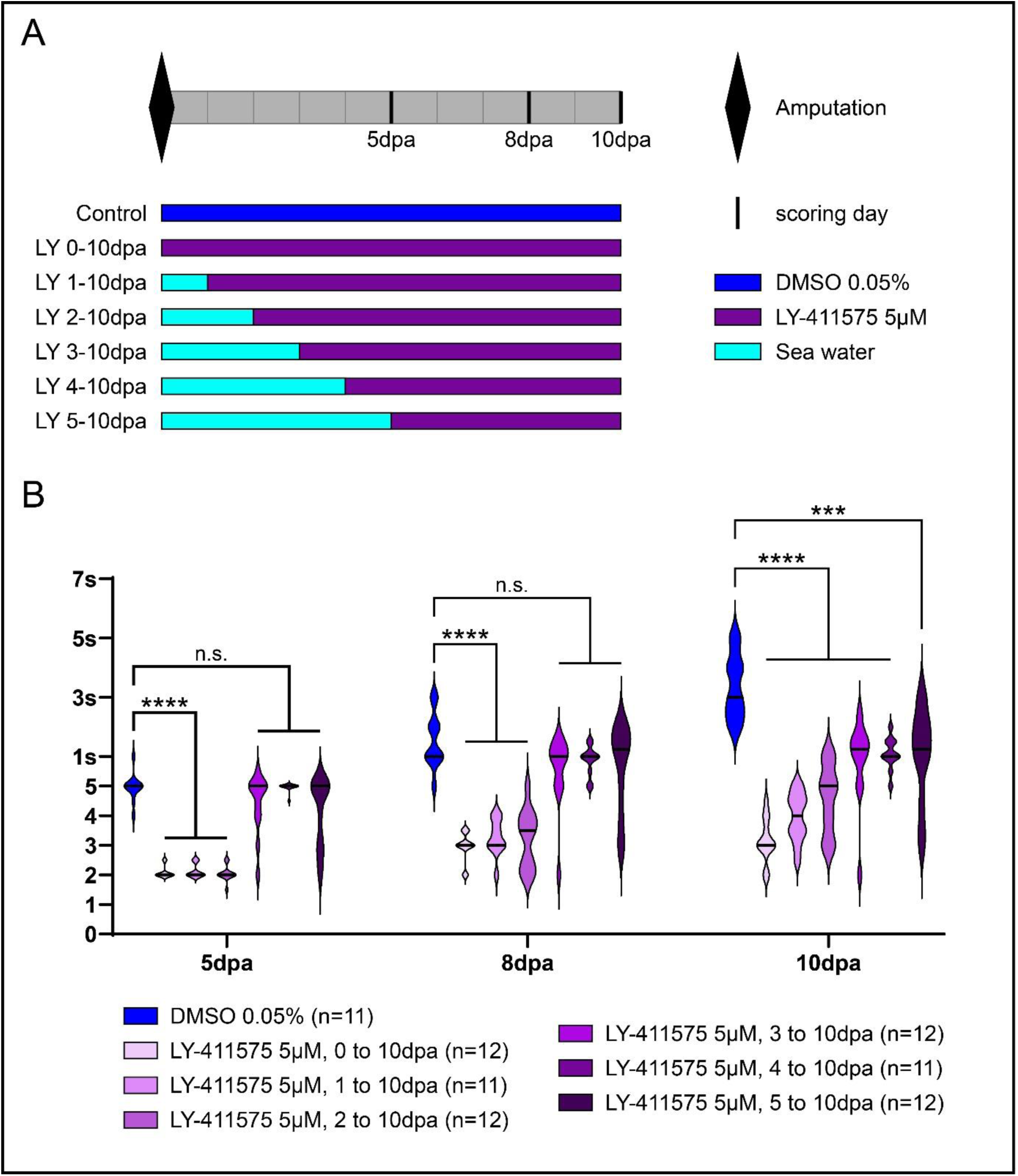
Morphological effects of different durations of Notch inhibition along posterior regeneration and posterior elongation. A) Schematic representation of the experiments: six durations of LY-411575 treatment were performed as well as a DMSO control. B) Violin plots representing the stages reached by each worm at 5, 8 and 10 dpa for each treatment. “n” represents the number of worms used per condition. Each bracket corresponds to a Mann-Whitney U test. n.s. p >0.05; * p<0.05; ** p<0.01; *** p<0.001; **** p<0.0001. dpa = day(s) post-amputation.

**Supplementary Figure 8:**
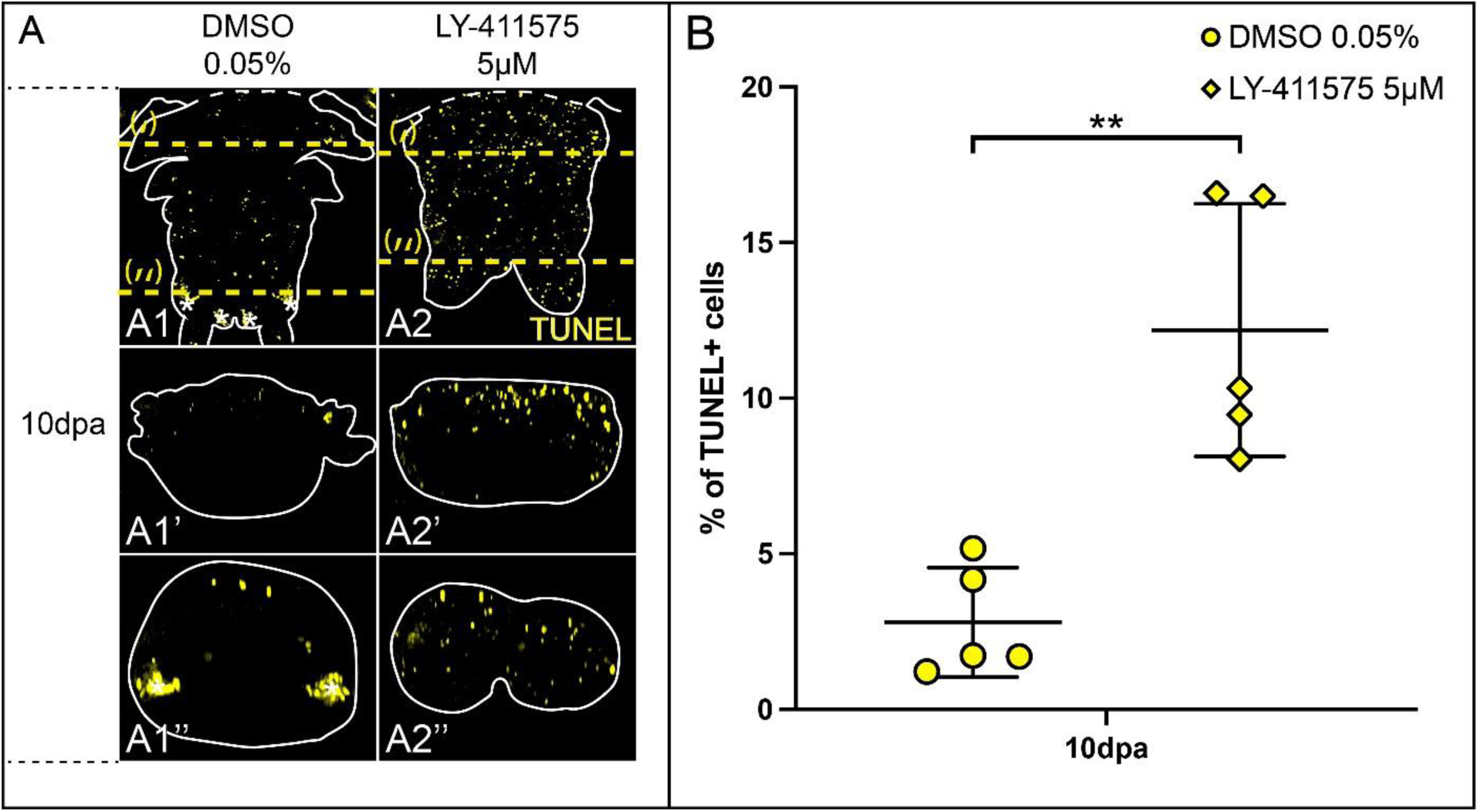
Notch inhibition during post-regeneration posterior elongation triggers apoptosis. A) TUNEL assay on whole-mount regenerated parts of LY-411575-treated worms from 3 dpa to 10 dpa and DMSO controls at 10 dpa. Ventral views are on the top and corresponding virtual transverse sections along the yellow dotted lines (‘ and ‘’, respectively) are at the bottom. Solid white lines delineate the outlines of the samples and white dashed lines correspond to the amputation planes. White asterisks = non-specific staining from parapodial glands. B) Proportions of TUNEL+ cells between LY-411575-treated worms from 3 dpa to 10 dpa and controls at 10 dpa. Each bracket corresponds to a Mann-Whitney U test. ** p<0.01. dpa = day(s) post-amputation.

**Supplementary Figure 9:**
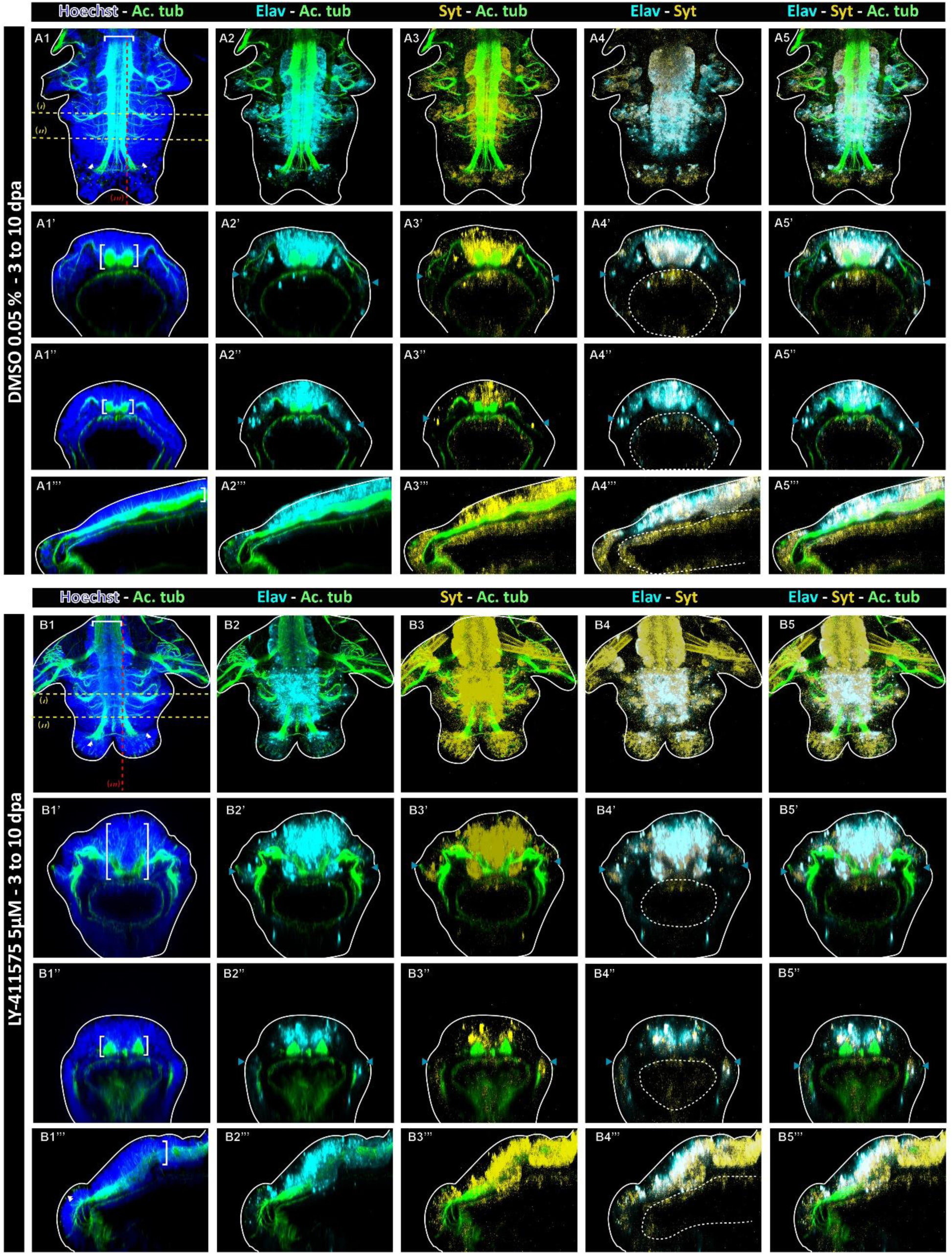
The neuronal marker *Syt* confirms the phenotype obtained upon Notch pathway inhibition during post-regeneration posterior elongation. Hybridization chain reactions (HCR) for *Elav* (cyan) and *Syt* (yellow) coupled with immunolabelling for acetylated tubulin (green) and nuclei staining with Hoechst (blue) for controls at 10 dpa (top) and LY-411575 treated regenerated parts from 3 dpa to 10 dpa (bottom). Ventral views are on top for each condition and corresponding virtual transverse sections (along (i) and (‘’) in yellow white) and sagittal section (along (iii) in red) are at the bottom. Solid white lines delineate the outlines of the sample, and white dotted lines delineate the gut. White brackets = ventral nerve chord; white double arrowheads = cirri nerves; blue arrowheads = PNS. dpa = day(s) post-amputation.

**Supplementary Figure 10:**
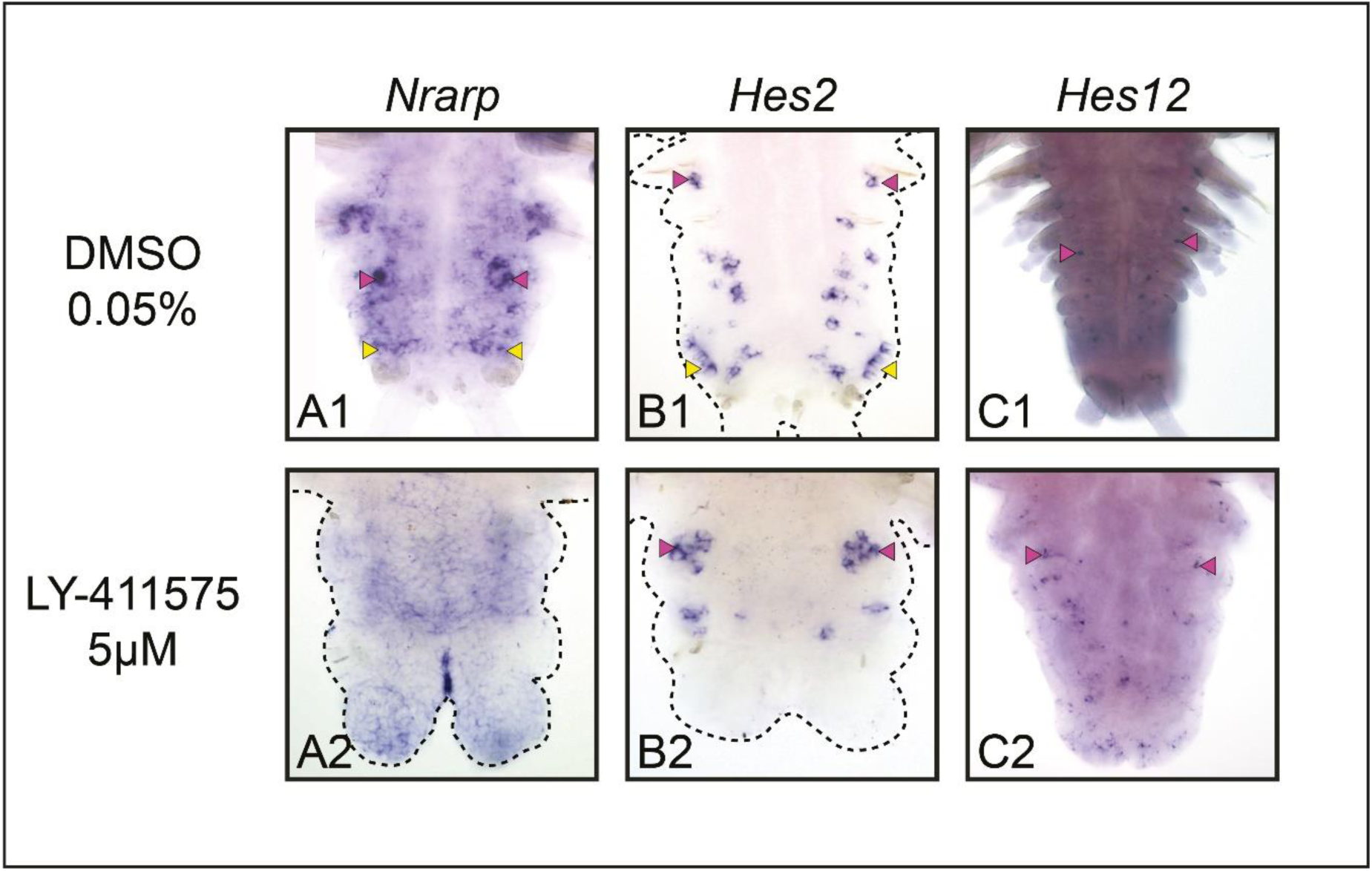
Notch inhibition during post-regeneration posterior elongation impairs chaetogenesis. Whole-mount *in situ* hybridizations for *Nrarp* and *Hes* genes expressed in chaetal sacs for LY-411575-treated worms from 3 dpa to 10 dpa and controls at 10 dpa. Ventral views. Yellow arrowheads = growth zone involved in posterior elongation of the animals [28]; pink arrowheads = chaetal sacs producing the parapodial bristles; dpa = day(s) post-amputation

**Supplementary Table 1: TMM expression values for Notch pathway components and members of the HES superfamily during posterior regeneration.**

Identifiers and TMM expression values for all investigated genes (11 members of the Notch pathway and 15 genes of the HES superfamily) for 6 stages of regeneration. RNA-seq data are from [35].

**Supplementary Table 2: Accession and detailed information about the RNA-seq data produced in this study.**

Detailed information for each dataset includes: (i) the name of the sample, (ii) the data type (short read, read length), (iii) the permanent links to data and accession numbers. Se= single end, rep= replicate

**Supplementary Table 3: List of differentially expressed genes (DEGs) between LY-411575 and DMSO conditions at 1 dpa.**

A gene is considered as differentially expressed with an FDR < 0.05 and a logFC > 1 or < −1. The table includes various annotations (BLAST) and GO terms. dpa= day post amputation

**Supplementary Table 4: List of differentially expressed genes (DEGs) between LY-411575 and DMSO conditions at 2 dpa.**

A gene is considered as differentially expressed with an FDR < 0.05 and a logFC > 1 or < −1. The table includes various annotations (BLAST) and GO terms. dpa= day post amputation

**Supplementary Table 5: TMM expression values for Notch pathway components and members of the HES superfamily for LY-411575 and DMSO conditions at 1 and 2 dpa.**

TMM expression values for all investigated genes (11 members of the Notch pathway and 15 genes of the HES superfamily) for all replicates during Notch inhibition at 1 and 2 dpa. Values for all replicated and mean are mentioned. Genes that are either downregulated or upregulated (at 1 or 2 dpa) are highlighted in orange and cyan, respectively. dpa= day post amputation

**Supplementary Table 6: Probes list for Hybridization Chain Reaction (HCR) experiments**

Sequences of the probes for the four genes investigated in this study.

## Acknowledgments

We are grateful to all past and present members of the Gazave lab for their support and suggestions on this study. We thank the Institute Jacques Monod facility staff for their help with the *Platynereis* culture. We acknowledge the ImagoSeine core facility of Institut Jacques Monod, member of France-BioImaging (ANR-10-INBS-04) and IBiSA, with the support of Labex “Who Am I”, Inserm Plan Cancer, Region Ile-de-France and Fondation Bettencourt Schueller.

## BIBLIOGRAPHY

1. Gazave, E., et al., Origin and evolution of the Notch signalling pathway: an overview from eukaryotic genomes. BMC Evol Biol, 2009. 9: p. 249.

2. Lv, Y., et al., Evolution and Function of the Notch Signaling Pathway: An Invertebrate Perspective. Int J Mol Sci, 2024. 25(6).

3. Bray, S.J., Notch signalling: a simple pathway becomes complex. Nat Rev Mol Cell Biol, 2006. 7(9): p. 678–89.

4. Henrique, D. and F. Schweisguth, Mechanisms of Notch signaling: a simple logic deployed in time and space. Development, 2019. 146(3).

5. Andersson, E.R., R. Sandberg, and U. Lendahl, Notch signaling: simplicity in design, versatility in function. Development, 2011. 138(17): p. 3593–612.

6. Hori, K., A. Sen, and S. Artavanis-Tsakonas, Notch signaling at a glance. J Cell Sci, 2013. 126(Pt 10): p. 2135–40.

7. Gazave, E. and E. Renard, Evolution of Notch Transmembrane Receptors, in Encyclopedia of Life Sciences (ELS). 2010, John Wiley & Sons: Ltd: Chichester.

8. Gazave, E., A. Guillou, and G. Balavoine, History of a prolific family: the Hes/Hey-related genes of the annelid Platynereis. Evodevo, 2014. 5: p. 29.

9. Kageyama, R., T. Ohtsuka, and T. Kobayashi, The Hes gene family: repressors and oscillators that orchestrate embryogenesis. Development, 2007. 134(7): p. 1243–51.

10. Cau, E. and P. Blader, Notch activity in the nervous system: to switch or not switch? Neural Dev, 2009. 4: p. 36.

11. Sjoqvist, M. and E.R. Andersson, Do as I say, Not(ch) as I do: Lateral control of cell fate. Dev Biol, 2019. 447(1): p. 58–70.

12. Bahrampour, S. and S. Thor, The Five Faces of Notch Signalling During Drosophila melanogaster Embryonic CNS Development. Adv Exp Med Biol, 2020. 1218: p. 39–58.

13. Sood, C., et al., Notch signaling regulates neural stem cell quiescence entry and exit in Drosophila. Development, 2022. 149(4).

14. Pierfelice, T., L. Alberi, and N. Gaiano, Notch in the vertebrate nervous system: an old dog with new tricks. Neuron, 2011. 69(5): p. 840–55.

15. Chouly, M. and L. Bally-Cuif, Generating neurons in the embryonic and adult brain: compared principles and mechanisms. C R Biol, 2024. 347: p. 199–221.

16. Kuzina, I., J.K. Song, and E. Giniger, How Notch establishes longitudinal axon connections between successive segments of the Drosophila CNS. Development, 2011. 138(9): p. 1839–49.

17. Kannan, R., et al., Tyrosine phosphorylation and proteolytic cleavage of Notch are required for non-canonical Notch/Abl signaling in Drosophila axon guidance. Development, 2018. 145(2).

18. Zhang, Y., et al., Notch-dependent binary fate choice regulates the Netrin pathway to control axon guidance of Drosophila visual projection neurons. Cell Rep, 2023. 42(3): p. 112143.

19. Aujla, P.K., et al., The Notch effector gene Hes1 regulates migration of hypothalamic neurons, neuropeptide content and axon targeting to the pituitary. Dev Biol, 2011. 353(1): p. 61–71.

20. Shi, M., et al., Forced notch signaling inhibits commissural axon outgrowth in the developing chick central nerve system. PLoS One, 2011. 6(1): p. e14570.

21. Lampada, A. and V. Taylor, Notch signaling as a master regulator of adult neurogenesis. Front Neurosci, 2023. 17: p. 1179011.

22. Morizet, D., et al., Reconstruction of macroglia and adult neurogenesis evolution through cross-species single-cell transcriptomic analyses. Nat Commun, 2024. 15(1): p. 3306.

23. Ozpolat, B.D., et al., The Nereid on the rise: Platynereis as a model system. Evodevo, 2021. 12(1): p. 10.

24. Schenkelaars, Q. and E. Gazave, The Annelid Platynereis dumerilii as an Experimental Model for Evo-Devo and Regeneration Studies, in Handbook of Marine Model Organisms in Experimental Biology-Established and Emerging, A. Boutet and B. Schierwater, Editors. 2021, CRC Press: Boca Raton. p. 23.

25. Denes, A.S., et al., Molecular architecture of annelid nerve cord supports common origin of nervous system centralization in bilateria. Cell, 2007. 129(2): p. 277–88.

26. Simionato, E., et al., atonal- and achaete-scute-related genes in the annelid Platynereis dumerilii: insights into the evolution of neural basic-Helix-Loop-Helix genes. BMC Evol Biol, 2008. 8: p. 170.

27. Gazave, E., Q.I.B. Lemaitre, and G. Balavoine, The Notch pathway in the annelid Platynereis: insights into chaetogenesis and neurogenesis processes. Open Biology, 2017. 7(2).

28. Gazave, E., et al., Posterior elongation in the annelid Platynereis dumerilii involves stem cells molecularly related to primordial germ cells. Dev Biol, 2013. 382(1): p. 246–67.

29. Planques, A., et al., Morphological, cellular and molecular characterization of posterior regeneration in the marine annelid Platynereis dumerilii. Dev Biol, 2019. 445(2): p. 189–210.

30. Poss, K.D., Advances in understanding tissue regenerative capacity and mechanisms in animals. Nat Rev Genet, 2010. 11(10): p. 710–22.

31. Bely, A.E. and K.G. Nyberg, Evolution of animal regeneration: re-emergence of a field. Trends Ecol Evol, 2010. 25(3): p. 161–70.

32. Bideau, L., et al., Animal regeneration in the era of transcriptomics. Cell Mol Life Sci, 2021. 78(8): p. 3941–3956.

33. Tiozzo, S. and R. Copley, Reconsidering regeneration in metazoans: an evo-devo approach. Frontiers in Ecology and Evolution, 2015. 3: p. 67.

34. Galliot, B. and L. Ghila, Cell plasticity in homeostasis and regeneration. Mol Reprod Dev, 2010. 77(10): p. 837–55.

35. Paré, L., et al., Transcriptomic landscape of posterior regeneration in the annelid Platynereis dumerilii. BMC Genomics, 2023. 24(583).

36. Kostyuchenko, R.P., et al., FoxA expression pattern in two polychaete species, Alitta virens and Platynereis dumerilii: Examination of the conserved key regulator of the gut development from cleavage through larval life, postlarval growth, and regeneration. Dev Dyn, 2019. 248(8): p. 728–743.

37. Prud’homme, B., et al., Arthropod-like expression patterns of engrailed and wingless in the annelid Platynereis dumerilii suggest a role in segment formation. Curr Biol, 2003. 13(21): p. 1876–81.

38. Vullien, A., et al., The Rich Evolutionary History of the Reactive Oxygen Species Metabolic Arsenal Shapes Its Mechanistic Plasticity at the Onset of Metazoan Regeneration. Mol Biol Evol, 2025. 42(1).

39. Vullien, A., et al., [A trio of mechanisms involved in regeneration initiation in animals]. Med Sci (Paris), 2021. 37(4): p. 349–358.

40. Poss, K.D. and E.M. Tanaka, Hallmarks of regeneration. Cell Stem Cell, 2024. 31(9): p. 1244–1261.

41. Demilly, A., et al., Coe genes are expressed in differentiating neurons in the central nervous system of protostomes. PLoS One, 2011. 6(6): p. e21213.

42. Kerner, P., et al., Orthologs of key vertebrate neural genes are expressed during neurogenesis in the annelid Platynereis dumerilii. Evol Dev, 2009. 11(5): p. 513–24.

43. Demilly, A., et al., Involvement of the Wnt/beta-catenin pathway in neurectoderm architecture in Platynereis dumerilii. Nat Commun, 2013. 4: p. 1915.

44. Zakrzewski, A.C., Molecular Characterization of Chaetae Formation in Annelida and Other Lophotrochozoa, in Fachbereich Biologie, Chemie und Pharmazie der Freien. 2011, Berlin.

45. Bray, S., Notch signalling in Drosophila: three ways to use a pathway. Semin Cell Dev Biol, 1998. 9(6): p. 591–7.

46. Iso, T., L. Kedes, and Y. Hamamori, HES and HERP families: multiple effectors of the Notch signaling pathway. J Cell Physiol, 2003. 194(3): p. 237–55.

47. Scimone, M.L., et al., Coordinated neuron-glia regeneration through Notch signaling in planarians. PLoS Genet, 2025. 21(1): p. e1011577.

48. Vanderhaeghen, P. and H.J. Cheng, Guidance molecules in axon pruning and cell death. Cold Spring Harb Perspect Biol, 2010. 2(6): p. a001859.

49. Dorresteijn, A.W., et al., Molecular specification of cell lines in the embryo of Platynereis (Annelida). Rouxs Arch Dev Biol, 1993. 202(5): p. 260–269.

50. Vervoort, M. and E. Gazave, Zoological and molecular methods to study Annelida regeneration using Platynereis dumerilii, in Methods in Molecular Biology, S.S. David Carroll, Editor. 2022, Springer.

51. Bideau, L., et al., Variations in cell plasticity and proliferation underlie distinct modes of regeneration along the antero-posterior axis in the annelid Platynereis. Development, 2024. 151(20).

52. Choi, H.M.T., et al., Third-generation in situ hybridization chain reaction: multiplexed, quantitative, sensitive, versatile, robust. Development, 2018. 145(12).

53. Kuehn, E., et al., Segment number threshold determines juvenile onset of germline cluster expansion in Platynereis dumerilii. J Exp Zool B Mol Dev Evol, 2022. 338(4): p. 225–240.

54. Golde, T.E., et al., gamma-Secretase inhibitors and modulators. Biochim Biophys Acta, 2013. 1828(12): p. 2898–907.

55. Grotek, B., D. Wehner, and G. Weidinger, Notch signaling coordinates cellular proliferation with differentiation during zebrafish fin regeneration. Development, 2013. 140(7): p. 1412–23.

56. Munch, J., A. Gonzalez-Rajal, and J.L. de la Pompa, Notch regulates blastema proliferation and prevents differentiation during adult zebrafish fin regeneration. Development, 2013. 140(7): p. 1402–11.

57. Munder, S., et al., Notch-signalling is required for head regeneration and tentacle patterning in Hydra. Dev Biol, 2013. 383(1): p. 146–57.

58. Hamada, M., et al., Evolution of the chordate regeneration blastema: Differential gene expression and conserved role of notch signaling during siphon regeneration in the ascidian Ciona. Dev Biol, 2015. 405(2): p. 304–15.

59. Gahan, J.M., et al., Functional studies on the role of Notch signaling in Hydractinia development. Dev Biol, 2017. 428(1): p. 224–231.

60. Mashanov, V., et al., Active Notch signaling is required for arm regeneration in a brittle star. PLoS One, 2020. 15(5): p. e0232981.

61. Dray, N., et al., Dynamic spatiotemporal coordination of neural stem cell fate decisions occurs through local feedback in the adult vertebrate brain. Cell Stem Cell, 2021. 28(8): p. 1457–1472 e12.

62. Haas, B.J., et al., De novo transcript sequence reconstruction from RNA-seq using the Trinity platform for reference generation and analysis. Nat Protoc, 2013. 8(8): p. 1494–512.

63. McCarthy, D.J., Y. Chen, and G.K. Smyth, Differential expression analysis of multifactor RNA-Seq experiments with respect to biological variation. Nucleic Acids Res, 2012. 40(10): p. 4288–97.

64. Conway, J.R., A. Lex, and N. Gehlenborg, UpSetR: an R package for the visualization of intersecting sets and their properties. Bioinformatics, 2017. 33(18): p. 2938–2940.

65. Blighe, K., S. Rana, and M. Lewis. EnhancedVolcano: publication-ready volcano plots with enhanced colouring and labeling. 2021.

66. Bryant, D.M., et al., A Tissue-Mapped Axolotl De Novo Transcriptome Enables Identification of Limb Regeneration Factors. Cell Rep, 2017. 18(3): p. 762–776.

67. Wu, T., et al., clusterProfiler 4.0: A universal enrichment tool for interpreting omics data. Innovation (Camb), 2021. 2(3): p. 100141.

